# Host Plant and Light Regime Shape Limited, Whole-Body Transcriptional Rhythmicity in the Bird Cherry-Oat Aphid (*Rhopalosiphum padi*)

**DOI:** 10.64898/2026.10.12.758664

**Authors:** Daniel Kunk, Vamsi Nalam

**Affiliations:** Cell and Molecular Biology Graduate Program, Colorado State University, Fort Collins, Colorado, USA; Department of Agricultural Biology, Colorado State University, Fort Collins, Colorado, USA

**Keywords:** circadian rhythm, diurnal rhythm, aphid, *Rhopalosiphum padi*, bird cherry-oat aphid, RNA-seq, dryR

## Abstract

**Background:** Daily cycles in environmental conditions shape the biology of plants and their insect herbivores. Because aphids feed nearly continuously from the phloem to sustain rapid population growth, they maintain prolonged, direct contact with their hosts, which exposes them to plant-derived daily rhythms in addition to those arising from the environment. We investigated whole-body transcriptional rhythmicity in the bird cherry-oat aphid (Rhopalosiphum padi) under light-dark (LD) and constant darkness (DD) conditions while aphids fed on either wheat or an artificial diet sachet, sampling transcript expression every 4 h over a 48 h window.

**Results:** Only a small fraction of the transcriptome in R. padi displayed 24 h rhythms, and this fraction declined further as light and host cues were removed. Unexpectedly, more transcripts were rhythmic in DD than in LD on wheat. This trend was not present in diet-fed aphids and was not consistently detected across rhythmic detection tools or significance thresholds. Transcripts that were rhythmic in aphids feeding on wheat hosts had higher amplitudes compared to those feeding on diet. Peak expression (i.e., phase) for rhythmic transcripts clustered near dawn and pre-dusk (zeitgeber time, ZT0 and ZT12), except for transcripts detected as rhythmic in darkness-only on wheat, which peaked approximately 4 h later. Rhythmic transcripts detected in this study were functionally enriched for processes relevant to plant-aphid interactions, including carbohydrate metabolism and detoxification. Subsequent promoter analysis found E-box motifs in both rhythmic and non-rhythmic transcripts, and de novo promoter motif discovery on rhythmic transcripts identified recurring Mad/SMAD- and PAR-bZIP-like motifs.

**Conclusions:** Whole-body transcriptional rhythmicity in R. padi is limited in scope and appears strongly dependent on environmental and host-derived cues. The candidate transcripts and regulatory pathways identified in this study provide clear targets for tissue-specific, functionally minded follow-up studies to investigate the extent and regulatory mechanisms of endogenous physiological and behavioral rhythms in R. padi.

## Background

Earth’s rotation generates predictable, daily cycles that shape the biology of plants and their insect herbivores, including aphids. Both aphids and plants have internal circadian clocks that anticipate and coordinate responses to these recurring cycles [1]. Beyond daily timing, both plants and aphids utilize their clocks’ light-entrainment pathways to detect changes in photoperiod and regulate seasonal processes such as host-plant transitions and sexual reproduction [2, 3]. In agricultural systems, understanding how plant and aphid clocks interact across both diurnal and seasonal timescales, and which downstream physiological and behavioral processes they regulate, could provide a foundation for pest management practices that optimize the timing of interventions to increase overall efficiency [4, 5].

Plants are notably rhythmic organisms that undergo daily variation in primary metabolism, photosynthesis, carbon allocation, solute transport, and defense [6–11]. To feed, aphids use their stylet mouthparts to navigate between plant cells, reach the phloem, and continuously ingest sugar-rich sap, maintaining prolonged, direct contact with host tissues [12]. This tight spatial coupling over extended durations amplifies the potential influence of rhythmic host-plant processes on the daily biology of aphids, as aphids experience environmental rhythms in addition to those of their host plants.

Aphids are also rhythmic organisms, with endogenous clocks that modulate diverse behavioral and physiological outputs [13–17]. Aphid lineages share most of the canonical clock genes with other insects, and the rhythmic accumulation and localization of these components in specialized regions of the aphid brain have been shown in numerous studies [18–22]. Interestingly, aphid clocks differ from the well-studied *Drosophila* model, containing duplications, deletions, and highly divergent sequences, suggesting potential adaptation to needs unique to aphids [3, 23–25]. Many of these clock genes encode transcription factors that generate transcriptional rhythms by directly regulating downstream genes via conserved motif sequences on their promoter regions. These genes include additional transcription factors and signaling molecules, whose products can either regulate the core clock directly or contribute to driving rhythmic physiological and behavioral outputs [26–29].

Our previous work in the bird cherry-oat aphid (*Rhopalosiphum padi*), a globally distributed pest of cereal crops and a vector of the barley yellow dwarf virus, found that approximately 10% of the whole-body transcriptome undergoes a diurnal rhythm (as detected by Cosinor; *p* < 0.05, without Benjamini-Hochberg false discovery rate (FDR) correction) when feeding on host plants [30]. However, it remains unclear to what extent these transcriptional rhythms are driven by rhythmic environmental and host cues rather than being endogenously maintained by the circadian clock. In addition, the regulatory mechanism of this transcriptional rhythmicity is poorly understood. The agricultural and economic relevance, in addition to the global distribution of *R. padi,* makes it a compelling system for studying the extent of daily regulation on aphid biology with the broader goal of identifying mechanisms or processes that could be leveraged for pest management [31–34]. For example, if aphid feeding behavior or host immune suppression follows predictable daily schedules, pesticide applications or resistance-promoting crop treatments timed to these windows could improve both efficacy and efficiency [4, 5].

In this study, we utilized a 48 h time-course RNA-seq experiment to dissect the contributions of rhythmic light-dark and temperature cycles, host-derived cues, and endogenous regulation to transcriptional rhythmicity in aphids. We sampled *R. padi* whole-body mRNA under light-dark (LD) and constant darkness (DD) conditions while aphids fed on either host plants (a dynamic, rhythmic diet source) or artificial diet (a stable, arrhythmic diet source). Specifically, we asked (i) to what extent transcriptional rhythms in *R. padi* persist under free-running (DD) versus light-entrained (LD) conditions, (ii) how feeding on the host plant versus artificial diet influences the abundance, strength, and organization of these rhythms, and (iii) which functional pathways and candidate genes exhibit rhythmic expression patterns and what regulatory motifs underlie these oscillations. Rhythmic transcripts were identified with dryR, complemented by rhythm detection with Cosinor and JTK_CYCLE [35, 36]. To interpret the biological significance of rhythmic classes detected by dryR, we performed functional enrichment analyses. We examined promoter regions for enriched transcription factor binding motifs, including directly scanning sequences for E-box elements, which are known targets of circadian clock transcription factors. This study provides an enhanced understanding of the prevalence and organization of rhythmic transcription in the presence and absence of host plant and environmental cues in *R. padi* and identifies transcript candidates and pathways that may be under direct circadian clock regulation.

## Methods

### Aphid Colony Maintenance and Plant Growth Conditions

*R. padi* colonies were maintained on three-week-old barley plants (*Hordeum vulgare* var. AC Metcalfe) and reared in plant growth tents in long-day conditions (16 h light-8 h dark). New colonies were started by placing 10-20 mixed-age individuals onto fresh barley plants every two weeks or when the colony became overcrowded. Plants were potted using PRO-MIX BX General Purpose Growing Medium (Premier Tech Horticulture, Quakertown, PA) in 6.35 x 6.35 x 7.62 cm plastic pots. In LD, light conditions were approximately 3,000 lux, and temperatures oscillated diurnally between 25 and 19 °C. In DD, environmental conditions remained constant at approximately 19 °C and 0 lux. Full environmental data for age synchronization, light entrainment, and sample collection windows on both wheat plants and artificial diet sachets are presented in Fig. S1.

### Artificial Diet Preparation and Feeding Sachet Assembly

Artificial diets were prepared using a modified Auclair formulation (see Table S1 for the complete ingredients list) [37]. Modifications included substituting outdated products for modern alternatives. Dry ingredients were weighed using a digital balance (for quantities > 10 mg) or an ultra-balance (model ME-5, Sartorius, Göttingen, Germany; for quantities ≤10 mg) and combined in a 2 L foil-wrapped Erlenmeyer flask. Deionized water and a magnetic stir bar were added, and the solution was stirred for 2 h at room temperature. The pH was adjusted to 7.6 with 5 M NaOH, and the diet was sterilized by filtration through a 0.2 μm bottle-top filter (Nalgene Rapid-Flow, Thermo Scientific, Waltham, MA, USA). Aliquots of 10 mL were dispensed into 15 mL conical tubes under aseptic conditions in a biosafety cabinet and immediately stored at -20 °C. All feeding sachets were additionally set up under aseptic conditions. Parafilm was stretched over the bottom rim of a 60 mm Petri dish. One mL of thawed diet was pipetted between the Parafilm and the Petri dish bottom. The Parafilm was pressed firmly along the dish wall to form an airtight seal. A nitrile O-ring (model 41UK25, Grainger, Lake Forest, IL, USA) was placed over the Parafilm, and after adding aphids using a paintbrush, the Petri dish lid was placed on top of the O-ring to prevent escape (Fig. S2A).

### Age Synchronization and Entrainment

To minimize developmental variation, aphids were age-synchronized within a 24 h window. *R. padi* adults (*n* = 10) were placed onto three-week-old wheat seedlings (*Triticum aestivum* var. Chinese Spring; Zadoks stage 1.3) at ZT8 (zeitgeber time; i.e., 8 h after lights-on) [38]. Aphids were confined to the 2^nd^ and 3^rd^ leaves using an aspirator and cylindrical cages constructed from 50 mL conical tubes. The conical tubes were cut at the base, hot-glued to 25.4 cm bamboo skewers, and sealed at both ends with cotton balls after adding aphids to allow air exchange while preventing escape (Fig. S2B). After 24 h (at ZT8), adults were counted and removed using an aspirator, leaving only nymphs born during the synchronization period. All cages were maintained in growth tents under LD conditions before entrainment and sampling.

### Wheat Experiments

Three days after adult removal, cages were randomly assigned to growth tents under either LD or DD conditions for 48 h before sample collection. Collection began at the onset of the first dark period following this entrainment period, referred to as ZT-8 (i.e., negative ZT8, or 8 h before lights on) for LD samples or CT-8 for DD samples (Circadian time; i.e., negative ZT8, or 8 h before subjective lights on) (Fig. S2C).

### Diet Experiments

For artificial diet experiments, age synchronization followed the same protocol as described above. However, nymphs were allowed to develop in their wheat cages for two additional days before transitioning to an artificial diet, to ensure that aphids reached adulthood. To begin entrainment, aphids were removed from their wheat cages using an aspirator and a paintbrush and pooled into 100 mm Petri dishes. Ten apterous adults were then added to artificial diet sachets and randomly assigned to LD or DD conditions just before lights-off for 48 h before collection began at ZT/CT-8 for samples in LD and DD, respectively. (Fig. S2D). In both light treatments, diet sachets were inverted, and a green sheet of craft paper was placed above the inverted dishes as a visual cue to feed and to shield the diet from direct light.

### Sample Collection

Aphid bodies were collected at 4 h intervals over 48 h, yielding 13 timepoints per treatment (*n* = 4 biological replicates per timepoint). Collection began at the onset of the first dark period following the 48 h entrainment, designated ZT-8 for LD samples or CT-8 for DD samples (Fig. S2C–E). To minimize light exposure, all samples collected at night or under DD were collected using a red LED headlight (model no. HDBIN32EB, Energizer, Missouri, USA). Five apterous adults were removed from each cage or sachet with a paintbrush, placed into a 1.7 mL microcentrifuge tube, and immediately frozen in liquid nitrogen (Fig. S2E). Samples were stored at -80 °C until RNA extraction.

### Survival and Performance Assays

To understand how aphid survival and performance differed among each light and host treatment, we conducted a separate 96 h performance assay under the same light and entrainment conditions described above. Ten age-synchronized apterous adults were assigned to either wheat cages or artificial diet sachets under each light regime (*n* = 3 wheat cages/diet sachets per condition). The number of surviving adults and nymphs was monitored for 96 h following entrainment utilizing a mechanical tally counter. For wheat samples, counts were taken at the end of the 96 h period to limit removing the wheat leaves from their cages, which risks significant aphid disruption and escape. For diet samples, counts were taken daily at ZT4 by gently flipping over the diet sachet and counting through the translucent Petri dish lid. At 72 and 96 h, honeydew droplets limited visibility through the Petri dish lid, and it was gently lifted off the O-ring while stabilizing the base of the dish to allow for an accurate count. Additionally, all daily counts taken for DD-treated diet-fed aphids were performed using a red LED headlamp. On artificial diet, daily survival and cumulative nymph counts were compared between LD and DD at each timepoint using a two-tailed t-test in R (v4.5.3) (hour 0 was excluded from testing due to zero variance) [39]. At the 96 h timepoint, adult survival and the total number of nymphs were compared across all four treatment conditions using a one-way ANOVA in R followed by a Tukey’s *post-hoc* test (Table S2). Compact letter displays were generated using the multcompView package (v0.1-11) [40].

### RNA Extraction

Sample IDs were randomized to generate lists of samples to extract that were balanced across timepoints and treatment conditions. RNA was then extracted in batches of 12 samples using the Quick-RNA Miniprep Kit following the manufacturer’s protocol (catalog no. R1055, Zymo Research, CA, USA). Briefly, the sample microcentrifuge tubes and plastic pestles were pre-chilled in liquid nitrogen for 30 s. Samples were then ground manually for 1 min. Lysis buffer (300 µL) was added immediately following grinding, and samples were thoroughly mixed with the pestle until no clumps of tissue remained visible. Lysates were further processed according to the manufacturer’s protocol, with on-column DNase I treatment and elution with 52 µL of DNase/RNase-free water. The quantity and purity of RNA were determined using a NanoDrop spectrophotometer (Thermo Scientific, MA, USA) with 2 µL of eluted RNA per sample.

### Quality Control, Sequencing, and RNA-Seq Analysis

RNA samples were submitted to the Novogene Corporation (Sacramento, CA, USA) in two batches for quality control, library preparation, and sequencing. Each batch contained 156 and 52 samples, respectively. The sequencing batch was used as a batch effect in downstream rhythmicity modeling with dryR. Sample integrity was assessed using an Agilent 5400 Fragment Analyzer (Agilent Technologies, CA, USA). All samples had RNA Integrity Numbers > 7.0 and passed quality control. Strand-specific mRNA libraries were prepared using poly(A) enrichment and paired-end (2x150 bp) sequenced on the NovaSeq X Plus platform (Illumina Inc., CA, USA) at a depth of approximately 30M raw read-pairs per sample. Raw sequencing reads were filtered and trimmed using fastp (v0.24.1) with default parameters [41]. Across all samples, > 99.5% of reads passed fastp quality filters. The mean (± standard deviation) number of reads per sample after filtering was 83.6 ± 14.7 M. The mean proportion of bases containing adapter sequences was 6.3 ± 1.2% per read. Furthermore, after quality filtering, more than 97% of bases were ≥ Q30, and the average duplication rate was 47.9 ± 7.3%, with an average GC content of 38.0 ± 0.5% per read. Transcript abundance was quantified using Salmon (v1.10.3) against a decoy-aware index generated from the *R. padi* reference genome and transcriptome (v4; GenBank assembly GCF_020882245.1) available from AphidBase (https://bipaa.genouest.org/is/aphidbase/) [42, 43]. The mean salmon mapping rate was 80.8 ± 1.2%, and the mean strand mapping bias was 0.50 ± 0.01. These metrics, generated by MutliQC (v1.28), indicate the libraries and quantifications were of high quality and suitable for downstream rhythmicity analysis (Table S3) [44]. Transcript-level quantifications were imported into R using the tximport package (v1.36.0) along with a custom TxDB object generated from the *R. padi* v4.0 genome annotation using the txdbmaker package (v1.4.2) [45].

### Rhythmicity Analysis

Rhythmicity analysis was performed using the dryR R package (v1.0.0; https://github.com/naef-lab/dryR), which enables modeling of both rhythmic and midpoint (i.e., mean) expression across treatments [35]. Because experiments on wheat and diet were conducted on different dates with distinct developmental timelines, these two datasets were analyzed separately to avoid confounding the date of experiment with host treatment effects (i.e., wheat vs diet). As a result, comparisons between host treatments are *ad hoc* rather than formal statistical statements. In the expression matrices, rows with a sum of 0, representing transcripts with no expression across any timepoint or replicate, were removed from all further analyses. The resulting wheat and diet expression matrices were analyzed using dryR’s dryseq() function with the treatment levels of LD and DD, a 24 h period, and the sequencing batch included as a batch effect. For treatments with two levels (i.e., LD and DD), there are five rhythmicity models (denoted with the prefix ‘R’) and two mean models (denoted with the prefix ‘M’) to which each transcript can be assigned (Fig. 1A). Model R1 includes arrhythmic transcripts with no detectable 24 h rhythm. Model R2 includes transcripts that are rhythmic only in LD (i.e., a diurnal rhythm). In contrast, model R3 includes transcripts that are rhythmic only in DD (i.e., dark-induced rhythm). Transcripts that maintain consistent rhythmicity across both LD and DD conditions are classified as model R4 (uniform circadian rhythm). If rhythmicity is maintained but shows changes in phase and/or amplitude between LD and DD conditions, these transcripts are assigned to model R5 (differential circadian rhythm). Independent of the rhythmicity model classification, transcripts can also be assigned to mean model M1, indicating shared midpoint expression between LD and DD, or to M2, indicating differential midpoint expression (Fig. 1A). The prefixes ‘Wh’ and ‘Ad’, added before a model label, represent its belonging to the wheat or artificial diet models, respectively. To ensure robust rhythmicity analysis, we filtered all transcripts assigned to rhythmicity models that had a Bayesian inference criterion weight (BICW) of less than 0.85, a log_2_ amplitude of less than 0.25, and a maximum Cook’s distance of 1 at a given timepoint. These transcripts were then assigned to a catch-all, filtered model designated as R0. Further, to limit the impact of large-amplitude transient spikes with poor expression support across timepoints, transcripts that were not expressed in at least three out of four replicates at a minimum of two timepoints were also assigned to model R0. To validate rhythmicity trends detected by dryR, Cosinor and JTK_CYCLE were run using the Nitecap web application, with DESeq2 - normalized reads generated by dryR used as the input matrices (https://nitecap.org/) [36].

**Fig. 1.**
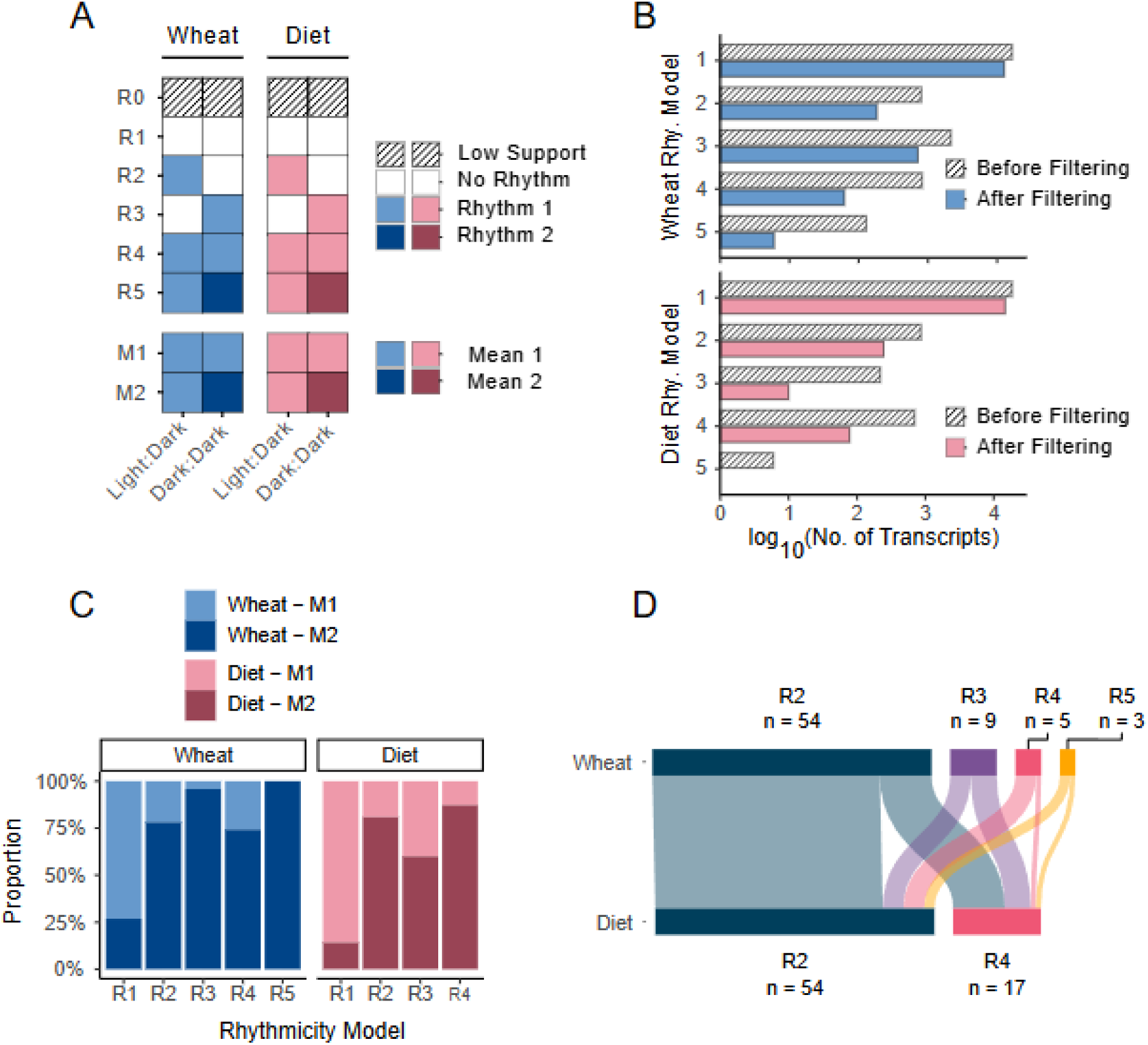
DryR model assignments and shared rhythmic transcripts across wheat and artificial diet. (A) Conceptual diagram showing how dryR assigns each transcript to rhythmicity (R1–R5) and mean (M1 and M2) models. The top row of striped squares represents a model containing transcripts that were removed from models R1–R5 due to low support for rhythmicity. White squares represent that no 24 h rhythm was present, while light-colored squares represent that the transcript had a 24 h rhythm in the corresponding light condition. Identical light squares within a row represent shared rhythm parameters (phase and amplitude) across light conditions. Rows with both light and dark squares indicate two distinct 24 h rhythms were detected with differing parameters (phase and/or amplitude). For mean models (M1 and M2), light squares indicate a transcript that had identical midpoint expression, whereas both light and dark squares indicate differing midpoint expression between the light conditions. All expressed transcripts in the wheat and diet experiments were assigned to one rhythmicity and one mean model. (B) Bar plot showing the log_10_-transformed counts of each dryR rhythmicity model before (striped) and after filtering (light blue/red) in the wheat (top) and artificial diet (bottom) experiments. (C) Stacked bar plots showing the proportion of transcripts assigned to dryR mean models (M1 and M2) within each rhythmicity model (R1–R5) for wheat (left, blue) and artificial diet (right, red). For each rhythmicity model on the x-axis, the corresponding bar segment length on the y-axis indicates the fraction of transcripts classified as M1 versus M2. (D) Sankey plot illustrating the relationship between transcripts detected as rhythmic (R2 -R5) in both the wheat and diet experiments. Solid rectangles represent dryR rhythmicity models on wheat (top) and diet (bottom). Flows indicate whether a transcript is assigned to the same or a different rhythmicity model across wheat and diet experiments. Sample sizes (n) label the number of transcripts in each rhythmic model that are also detected as rhythmic in the other experiment, whether assigned to the same or a different model.

### Functional Enrichment Analysis

For each rhythmic model identified by dryR, the corresponding transcript IDs were imported into OmicsBox for functional annotation and enrichment analysis (v3.5; BioBam Bioinformatics, Valencia, Spain). GO term and KEGG pathway enrichment using a two-tailed Fisher’s exact test was performed separately for all rhythmic models. All expressed transcripts modeled by dryR were used as the respective reference universe sets. Thus, GO terms and KEGG pathways enriched in each rhythmicity model are relative to the total set of expressed transcripts in the respective host treatment. Fold enrichment was calculated as:

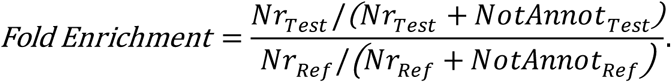

Here, *NrTest* is the number of rhythmic transcripts in a dryR model that were annotated to a particular term or pathway. Conversely, *NotAnnotTest* is the number of transcripts in the model that were not annotated to that term. *NrRef* and *NotAnnotRef* are the corresponding counts in the reference universe set.

### Promoter Motif Discovery and Enrichment Analysis

For promoter sequence analysis, the regions -1500 bp upstream and +150 bp downstream of the transcription start site (TSS) were acquired for all genes using the GenomicFeatures R package (v1.58.0) [46]. Fasta files containing the promoter regions that correspond to the transcripts in each dryR rhythmicity model were saved and used as input to the MEME suite of tools. For *de novo* motif enrichment analysis, the XSTREME (v5.5.8) tool was used to search against the JASPAR non-redundant core insect motif database (2022) with default settings [47]. The promoter sequences corresponding to transcripts in the dryR model R1 were used as control sequences. Thus, enriched motifs in each dryR rhythmicity model are relative to motifs in promoter regions of non-rhythmic transcripts (R1). Additionally, the FIMO tool with default settings was used to scan the promoters corresponding to the transcripts in each dryR rhythmicity model for E-box motifs [48]. The motifs scanned with FIMO included the canonical E-box motif CACGTG (JASPAR MA2301 and MA2305) and a universal E-box sequence of CANNTG.

## Results

### Adult Survival is Comparable Across Host and Light Treatments, While Reproductive Performance is Highest on Wheat In LD

To determine whether feeding on dark-stressed plants or artificial diet impacted aphid performance, we monitored survival and reproduction in all four light and host treatment combinations across 96 h (Fig. S3; Table S2). On artificial diet sachets, the number of surviving adults and the cumulative number of nymphs did not differ significantly between LD and DD at any timepoint (t-tests, all *p* > 0.05; Fig. S3A, B).

At 96 h, total surviving adults did not differ significantly across the four treatment groups (one-way ANOVA, *F*(3,8) = 1.6, *p* = 0.27; Fig. S3C). Total nymph production differed significantly among the four treatment groups, with aphids feeding on wheat under LD conditions producing significantly more nymphs than all other treatments (one-way ANOVA, *F*(3,8) = 32.0, *p* = 8.3x10^-5^; Fig. S3D). Removing light cues reduced nymph production more strongly on wheat than on artificial diet, with aphids feeding on wheat producing significantly fewer nymphs under DD (Tukey’s HSD, p = 0.033); however, nymph production did not differ significantly between wheat-fed aphids in DD and diet-fed aphids in LD (*p* = 0.283). Further, nymph production on diet did not differ significantly between LD and DD (*p* > 0.05), consistent with the daily comparisons reported above. Together, these results indicate that the host treatments used in our experiment supported adult aphid survival throughout the sampling window.

### Rhythmic Transcription is Limited and Influenced by Host Plant and Light Regime

Rhythmicity was evaluated across the expressed transcriptome, which comprised 96.2% and 96.1% of all annotated transcripts in the wheat and artificial diet experiments, respectively. In both host treatments, only a small subset of expressed transcripts showed a 24 h rhythm in either light regime as detected by dryR. On wheat, 4.7% of expressed transcripts (964 transcripts) cycled in at least one light condition; the remainder showed no detectable rhythm (12,633 transcripts) or was excluded for low model support (6,710 transcripts). Most rhythmic transcripts on wheat were specific to a single light regime: 183 cycled only under LD conditions (i.e., diurnal, WhR2) and 713 cycled only under constant darkness (i.e., dark-induced, WhR3). A smaller set of 68 transcripts were rhythmic under both LD and DD conditions; of these, 62 kept the same phase and amplitude conditions (i.e., shared- or circadian rhythm, WhR4), while 6 shifted in phase and/or amplitude between LD and DD (i.e., differential circadian rhythm, WhR5; Fig. 1B, top; Table S4).

Rhythmic transcription was reduced approximately threefold in aphids feeding on artificial diet: only 1.6% of expressed transcripts (333 transcripts) cycled in at least one light condition, while 14,644 transcripts were arrhythmic and 5,296 were excluded for low support. As seen in wheat, diurnal-only rhythmicity was the most common form (245 transcripts, AdR2), with a smaller group maintaining rhythmicity across LD and DD (78 transcripts, AdR4). Interestingly, darkness-induced rhythmicity was almost absent in diet-fed aphids as compared to their wheat counterparts (10 transcripts, AdR3). No transcripts showed differential circadian rhythms for diet-fed aphids after filtering for model support (AdR5; Fig. 1B, bottom; Table S4). A notable asymmetry was observed between feeding treatments in DD conditions, with high levels of rhythmic transcription in aphids feeding on wheat but little to no rhythmicity in aphids feeding on artificial diet.

Independent of whether a transcript was rhythmic, its average expression across the sampling window could also differ between LD and DD (Fig. 1A). Most arrhythmic transcripts showed similar average expression across light regimes in both host treatments (WhM1, AdM1), although roughly a quarter displayed differential midpoint expression on wheat and diet alike (WhM2, AdM2). Rhythmic transcripts, in contrast, were far more likely to show this shift, with the majority of transcripts in every rhythmic class, for both wheat- and diet-fed aphids, differing in midpoint expression between LD and DD (Fig. 1C; Table S4).

These rhythmicity classes were broadly, but not consistently, supported by two different rhythm-detection approaches: Cosinor and JTK_CYCLE. Agreement across all three approaches was strongest under LD conditions. In contrast, rhythmicity detected under DD, including the highly enriched class on wheat (WhR3), was less consistent across methods and more sensitive to *p*-value correction (Fig. S4; Table S5; Supplemental Note S1). This pattern suggests that the DD-associated signal on wheat, while robustly detected in the primary dryR modeling framework used throughout this study, has a comparatively weak or noisy oscillatory structure that warrants cautious interpretation.

### Rhythmicity is Largely Host-Dependent, with Diurnal Rhythms the Most Conserved on Artificial Diet

Comparing the rhythmic classifications between wheat and diet-fed aphids showed that rhythmicity was largely host-specific (Fig. 1D; Table S6). Few transcripts that were non-rhythmic on wheat became rhythmic on diet (63 transcripts), whereas substantially more transcripts that were non-rhythmic on diet were rhythmic on wheat (450 transcripts), most within the dark-induced classification (WhR3). Only 71 transcripts were rhythmic in both wheat- and diet-fed aphids. Of the 183 diurnally rhythmic transcripts identified on wheat (WhR2), 54 retained rhythmicity on diet, 44 remained diurnally rhythmic (AdR2), and 10 shifted to a shared LD/DD rhythm (AdR4). In contrast, dark-induced (WhR3) and differentially rhythmic (WhR5) transcripts on wheat showed little conservation and were largely lost or reclassified on artificial diet.

Two transcripts stood out by retaining rhythmic behavior across both host and light treatments and are strong candidates for direct, endogenous circadian regulation. A putative polycomb ortholog (XM_060995664.1) was the only transcript with a shared LD/DD rhythm on both wheat and diet (WhR4/AdR4); in *Drosophila*, polycomb was shown to regulate circadian rhythms and is required for normal sleep-wake cycling [49, 50]. Additionally, a putative neprilysin-2 ortholog (XM_060991018.1), which belongs to a family of peptidases implicated in feeding behavior and numerous other neurological processes, was rhythmic in LD and DD conditions on both hosts, though with a shifted phase between LD and DD on wheat (WhR5/AdR4) [51–53].

### Rhythmic Transcripts Show a Bimodal, Dawn and Pre-Dusk Phase Organization

Across both host treatments, diurnally rhythmic transcripts (WhR2/AdR2) exhibited peak expression (i.e., phase) as a bimodal distribution, with two distinct clusters at ZT0 (dawn) and ZT12 (pre-dusk) (Fig. 2; Table S4).

**Fig. 2.**
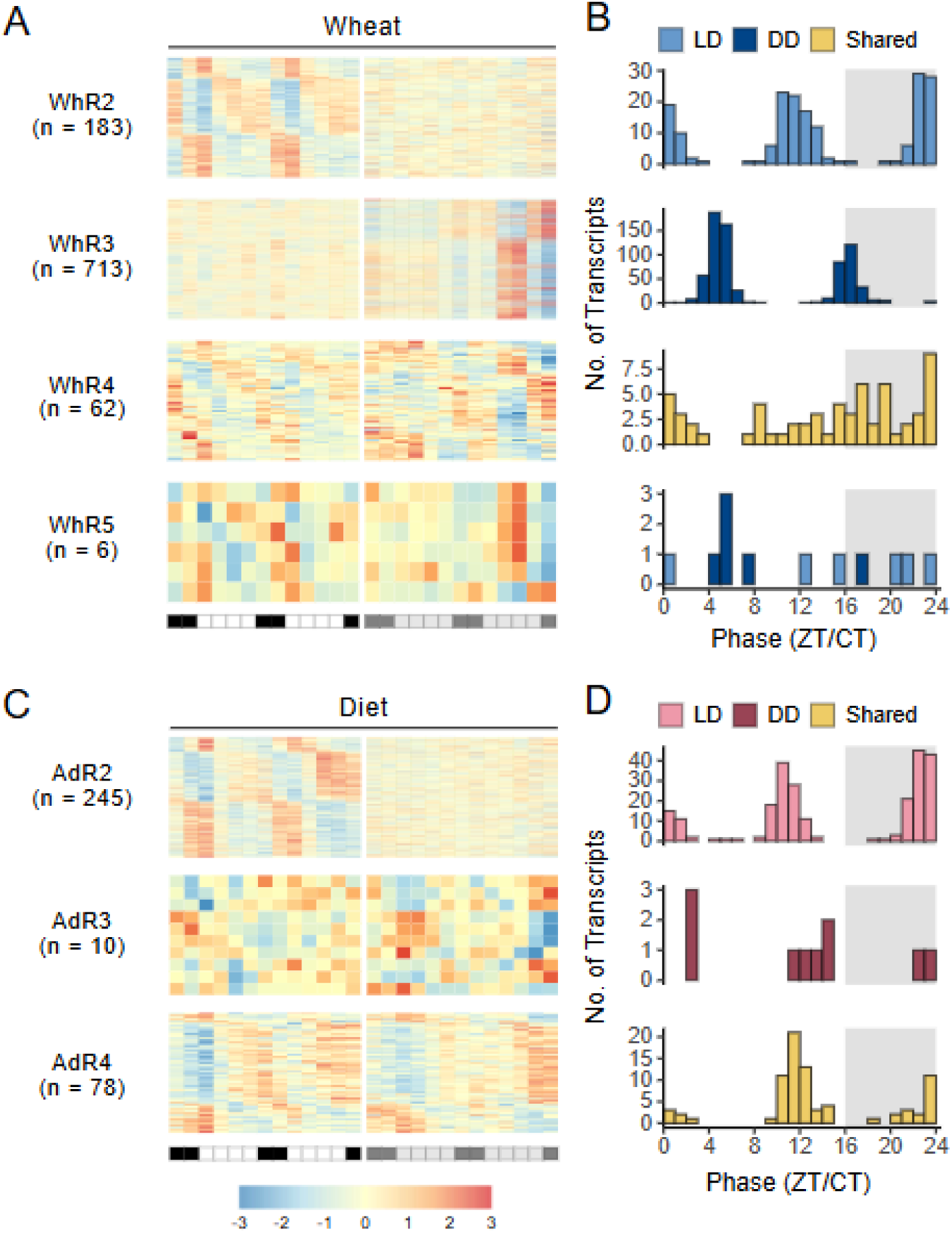
Rhythmic expression patterns and phase distributions across wheat and diet. Multi-panel plots showing transcript expression patterns and phase distributions for dryR rhythmicity models (R2-R5) in wheat (A-B) and artificial diet (C-D), with each row corresponding to a single rhythmicity model across the paired panels. Heatmaps (A, C) display the z-scored VST-normalized expression of transcripts assigned to each model. Heatmap columns represent timepoints and rows represent transcripts, and the timeline bar indicates time of day (white: day; black: night; gray: subjective day/night). For each model, the left and right heatmap panels show expression profiles under LD and DD, respectively. Sample size (n) indicates the number of transcripts assigned to each model after filtering. Histograms (B, D) show phase distributions for the same model as the corresponding heatmap, plotted in zeitgeber time (ZT) for LD and circadian time (CT) for DD.

Dark-induced transcripts on wheat (WhR3) also showed this bimodal phase pattern but roughly 4 h later into the day, clustering near CT4 and CT16 (Fig. 2A, B). This delay was not shared by the much smaller dark-induced transcripts classified on diet-fed aphids (AdR3), whose phases instead remained close to the LD-like dawn and pre-dusk phase pattern (Fig. 2C, D), reinforcing that the delayed DD phase on wheat may likely reflect aphids responding to a lagging, plant-derived signal rather than a direct impact of constant darkness itself.

Shared-rhythmic transcripts between LD and DD (WhR4/AdR4) showed similar overall oscillatory behavior in both host treatments, although their phase distribution slightly differed. On wheat, these transcripts had phase distributions across a broader spectrum of the day, whereas on diet the phases remained bimodal, again clustering near dawn and pre-dusk. Differentially rhythmic transcripts on wheat (WhR5) shifted from later phases in LD to earlier, more tightly clustered phases in DD, converging near ZT/CT4 -8 and ZT/CT16 (Fig. S5; Table S4).

Phases estimated independently by JTK_CYCLE followed the same overall pattern, with peak expression concentrated near dawn and pre-dusk across host and light treatments (Fig. S6A; Table S5).

### Amplitude of Rhythmic Transcripts is Highest on Wheat and Under Light-Dark Conditions

Amplitude distributions were right-skewed across all treatments and rhythmicity classifications, with most rhythmic transcripts showing a log_2_ fold-change (log_2_FC) amplitude below 2.25 (Fig. 3; Table S4).

**Fig. 3.**
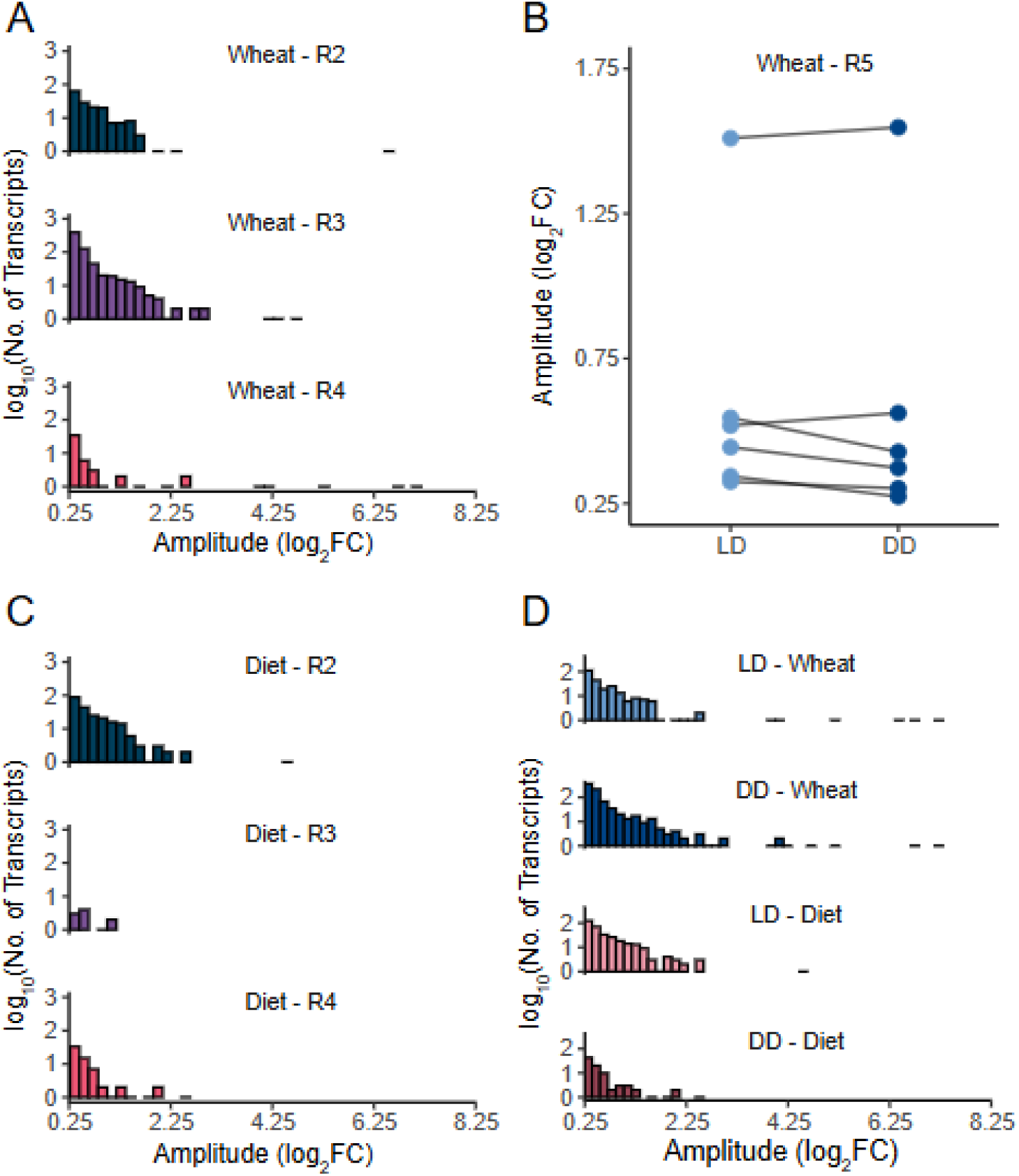
Amplitude distributions of rhythmic transcripts across dryR models and treatments. (A) Histograms showing amplitude distributions for rhythmic transcripts assigned to dryR models R2-R4 in wheat. The y-axis represents the log_10_-transformed number of transcripts per bar, while the x-axis represents the log₂ fold-change amplitude. (B) Paired dumbbell plot showing log₂ fold-change amplitude for transcripts assigned to model R5 in wheat in LD and DD. Dots indicate the corresponding amplitude, and connecting lines indicate the difference in amplitude between LD and DD. (C) Histograms showing amplitude distributions for rhythmic transcripts assigned to dryR models R2 -R4 in artificial diet. (D) Amplitude distributions for rhythmic transcripts when grouped by light condition (LD vs DD) and source (wheat vs diet).

On wheat, diurnal and dark-induced transcripts (WhR2 and WhR3, respectively) had comparable median amplitudes (0.49 ± 0.58 and 0.40 ± 0.46 log_2_FC, respectively), whereas the shared-rhythm (WhR4) class included a longer tail of high-amplitude transcripts but had a higher density of low-amplitude transcripts, resulting in a slightly lower median amplitude (0.37 ± 1.48 log_2_FC). The small set of differentially rhythmic transcripts on wheat (WhR5) had uniformly moderate median amplitudes that were slightly lower in DD than LD (0.40 ± 0.48 and 0.48 ± 0.45 log_2_FC, respectively). On artificial diet, diurnal and shared-rhythmic transcripts (AdR2 and AdR4, respectively) showed similarly right-skewed median amplitudes (0.49 ± 0.68 and 0.37 ± 0.45 log_2_FC, respectively), as did the small, dark-induced (AdR3) rhythmic class (0.52 ± 0.28 log_2_FC; Fig. 3A, B, C).

When grouped by light and host treatment, rhythmic transcripts consistently reached higher median amplitudes under LD than DD, and on wheat than on diet. Median amplitude was 0.45 ± 0.90 log_2_FC on wheat in LD versus 0.40 ± 0.61 log_2_FC in DD, and 0.47 ± 0.49 log_2_FC on diet in LD versus 0.41 ± 0.43 log_2_FC in DD (Fig. 3D). Amplitudes estimated independently by JTK_CYCLE showed the same right-skewed distributions and relative differences between treatments (Fig. S6B; Table S5). Together, rhythmic transcripts oscillate with consistently larger amplitudes in the presence of light and the host plant.

### Rhythmic Transcripts are Functionally Enriched for RNA-Processing and Possible Plant-Aphid Interaction Pathways

On wheat, diurnally rhythmic transcripts (WhR2) were nominally enriched (*p* < 0.05 but not passing the FDR correction threshold of *q* < 0.05) for broad regulatory functions such as regulation of macromolecule and primary metabolic processes. These enriched GO terms were driven by putative serine/threonine kinases, transcription factors, core clock components (casein kinases, shaggy, cycle, etc.; Fig. S7A; Table S7). Further, nominally enriched KEGG pathways in this class centered on signaling and nucleotide metabolism, including hedgehog and IL-17 signaling and pyrimidine metabolism (Fig. S7B; Table S8).

Dark-induced transcripts on wheat (WhR3) showed the strongest functional enrichment of any signal class, with 33 GO terms significant after FDR correction (*q* < 0.05). These terms were mainly centered around protein folding and processing, such as organelle lumen, unfolded protein binding, and protein folding. These enriched terms were driven by putative T-complex chaperonin subunits, prefoldin components, calreticulin, and multiple heat shock proteins (Fig. S7A; Table S7). Nominally enriched KEGG pathways pointed to amino acid and carbon metabolism, including glycine/serine/threonine metabolism, carbon fixation, and tyrosine metabolism, which included putative pyruvate carboxylase, malate dehydrogenase, aldehyde dehydrogenase, and serine hydroxymethyltransferase orthologs (Fig. S7B; Table S8). Together, these enrichments suggest that dark-induced transcripts in wheat are heavily biased toward stress, as well as carbohydrate and protein metabolic pathways. Because light serves as both the plant’s primary energy source and a daily cue, its removal likely triggers a cascading metabolic stress from host to aphid. Interestingly, these transcripts becoming rhythmically expressed, yet phase shifted relative to transcripts in other rhythmic classes, may suggest that the metabolic state of the host plant and the aphid can reorganize transcriptional rhythmicity.

Transcripts with shared rhythmicity between LD and DD on wheat (WhR4) were significantly enriched (*q* < 0.05) for six GO terms related to muscle structure and contraction, such as troponin complex, myofilaments, and muscle contraction, driven by putative troponin and myosin isoforms. Nominally enriched KEGG pathways spanned glycan biosynthesis (O-glycan, N-glycan) and hormone and retinol metabolism, largely driven by putative UDP-glycosyltransferases UGT4 and UGT5 (Fig. S7A, B; Table S7, S8).

The small set of differentially rhythmic transcripts in wheat (WhR5) was only nominally enriched with GO terms such as exon-exon junction complex and sexual reproduction-related terms. These results should be interpreted cautiously, as a single transcript drove each term; specifically, two putative isoforms of partner of Y14 and mago in addition to neprilysin-2.

On artificial diet, diurnally rhythmic transcripts (AdR2) were the only class with GO terms surviving FDR correction (*q* < 0.05). Specifically, a set of transcription start site selection-related terms was driven by putative serine/arginine-rich splicing factor 1 (RpSrsf1) isoforms and RNA-binding protein 1-like. An additional 68 terms were nominally enriched around RNA metabolism and biosynthesis, driven by numerous RNA-binding proteins, kinases, and splicing factors (Fig. S8A; Table S7). Nominally enriched KEGG pathways reflected RNA processing and signaling, driven by overlapping sets of splicing and translation-initiation factors and numerous zinc-finger proteins (Fig. S8B; Table S8).

Dark-induced transcripts (AdR3) on diet were only nominally enriched, mostly by single transcripts, for cytoskeletal, metabolic, and neuronal terms such as actin-myosin filament sliding, pentose phosphate shunt, and nucleotide transport. These terms were driven respectively by putative unconventional myosin-IXAa-like, ribose-5-phosphate isomerase, and mitochondrial folate transporter orthologs; the single enriched KEGG pathway, motor proteins, was driven by the myosin-IXAa-like ortholog (Fig. S8A, B; Table S7, S8).

Transcripts with shared rhythms between LD and DD on diet (AdR4) were nominally enriched for 272 GO terms, including carbohydrate metabolic process, which was driven by putative phosphoglucomutase, glycogen debranching enzyme, insulin-degrading enzyme, phosphorylase b kinase gamma, and the salivary effector ortholog xylulose reductase-like [54]. Several developmental-related GO terms were also nominally enriched, including wing vein morphogenesis and endothelial tube morphogenesis. KEGG pathways additionally emphasized carbohydrate metabolism and detoxification, including the terms pentose phosphate pathway and starch and sucrose metabolism (Fig. S8A, B; Table S7, S8).

### E-Box Motifs are Present in Both Rhythmic and Non-Rhythmic Transcripts

Scanning promoter regions (-1500 to +150 bp relative to the TSS) for canonical E-box motifs (CACGTG/CANNTG) showed that E-boxes were present in roughly 10% of promoters regardless of rhythmicity status, with only slight variation by rhythmic class or host treatment. Specifically, E-box frequency was similar between arrhythmic and rhythmic classes on wheat (9.8-16.0% across rhythmic classes vs. 10.6% in non-rhythmic transcripts). Still, it was more variable on diet, ranging from 5.1% (shared-rhythmic transcripts, AdR4) up to 20.0% (dark-induced transcripts, AdR3) against a 10.4% baseline in arrhythmic transcripts (AdR1) (Fig. 4; Table S9). E-box positions were widely distributed across the promoter region, with fewer motifs occurring past the TSS depending on rhythmicity class and host treatment (Fig. 5).

**Fig. 4.**
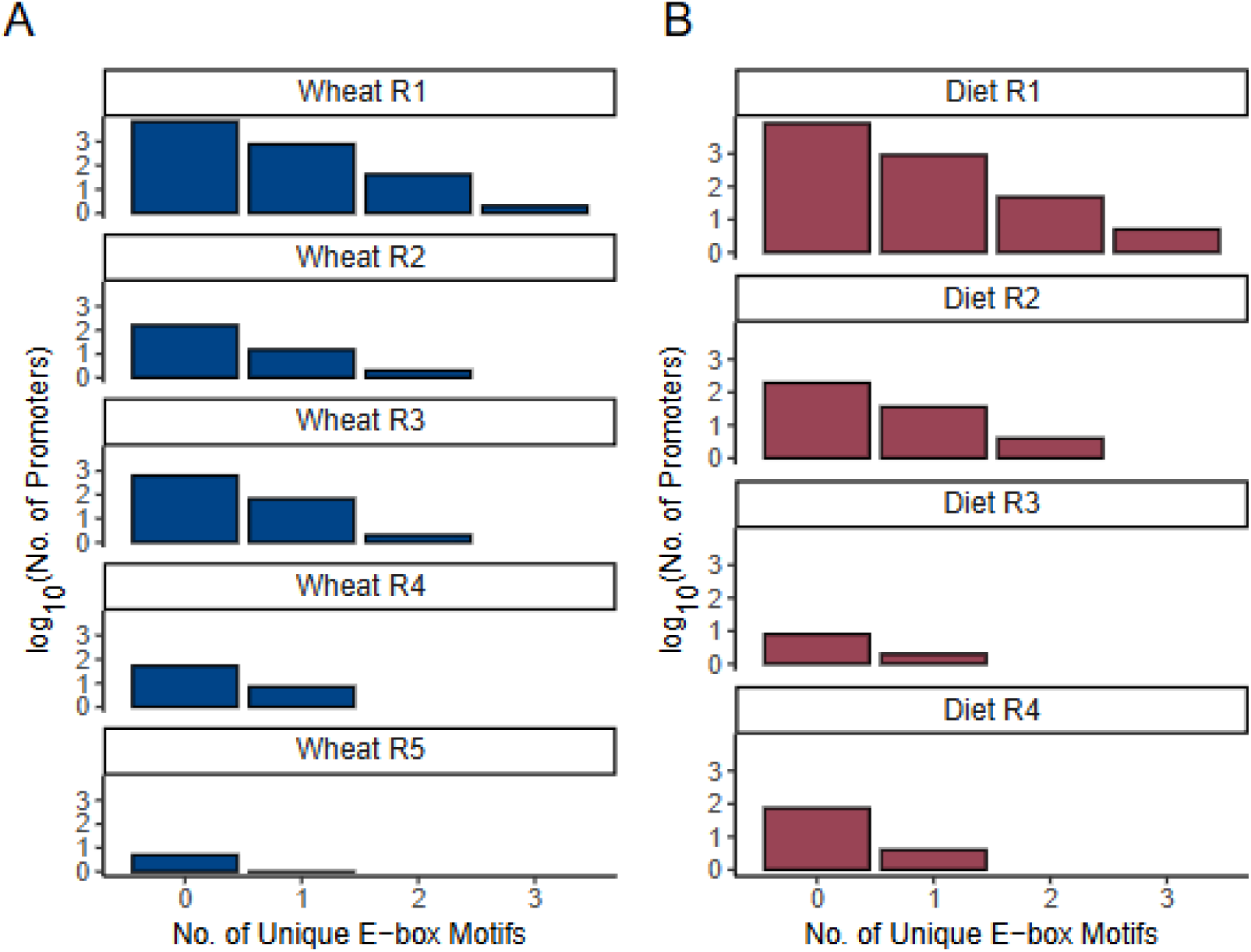
Distribution of unique E-box motifs per promoter in dryR rhythmicity models. Histograms showing the distribution of the number of unique E-box motifs detected per promoter region (−1500 bp upstream to +150 bp downstream of the TSS) for transcripts assigned to dryR rhythmicity models in wheat (A) and artificial diet (B). The y-axis indicates the log₁₀ number of promoters with a given motif count. Unique motifs were defined by collapsing overlapping hits from the canonical E-box (CACGTG) and universal E-box (CANNTG) motifs, such that matches at the same genomic coordinates were counted only once.

**Fig. 5.**
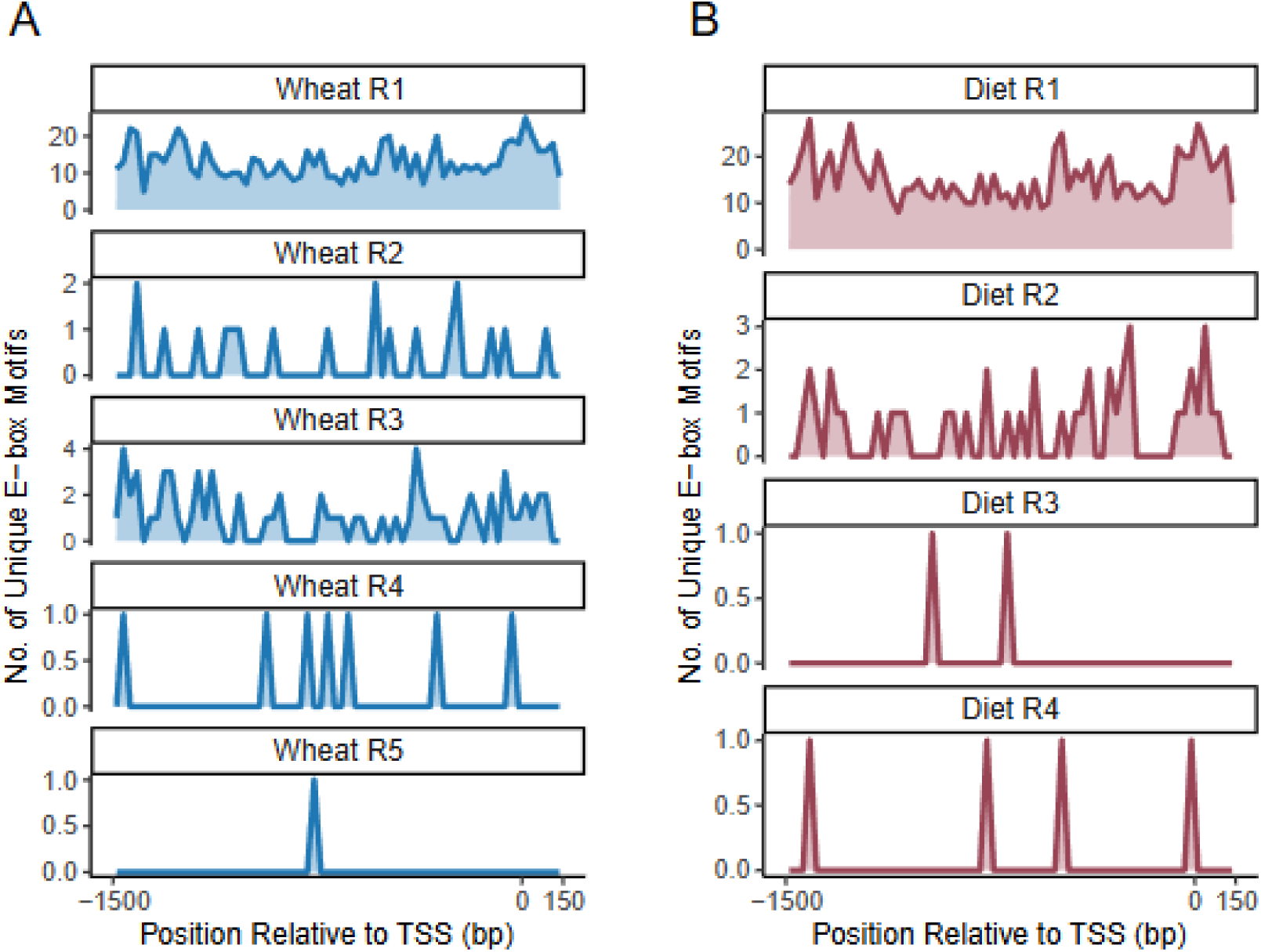
Positional distribution of unique E-box motifs relative to the transcription start site across dryR models. Positional distributions of unique E-box motifs relative to the transcription start site (TSS) for transcripts assigned to dryR rhythmicity models in wheat (A) and artificial diet (B). Curves show the number of unique E-box motifs detected at each position within promoter regions spanning −1500 bp upstream to +150 bp downstream of the TSS (0 bp). Motif counts are “unique” because overlapping matches to the canonical E-box (CACGTG) and the universal E-box (CANNTG) motifs at the same genomic coordinates were collapsed and counted only once.

*De novo* motif discovery identified additional regulatory signatures enriched specifically among rhythmic transcript promoters relative to arrhythmic promoters (E < 0.05; Fig. S9–12; Table S10). Across most rhythmicity classes in both host treatments, the dominant signal was a recurring GC-rich motif resembling Mad/SMAD binding sites, alongside bZIP and C2H2 zinc-finger (including Krüppel-like) motifs; the shared-rhythm class (Wh/AdR4) additionally showed enrichment for hair-related basic helix-loop-helix-like (bHLH) motifs in both wheat and diet. Diurnally rhythmic transcripts on wheat (WhR2) additionally contained a distinct, and potentially clock-relevant Crg-1 motif. In contrast, the differentially rhythmic class (WhR5) departed from the broader Mad-dominated pattern and instead contained low-occupancy and AT-rich bZIP, bHLH (SREBP-like), and TATA-like motifs. Further, diurnally rhythmic transcripts on diet (AdR2) carried additional bZIP (erect wing-like) and C2H2 (squeeze-like) motifs, while dark-induced transcripts on diet (AdR3) showed a more heterogeneous and poorly supported motif background dominated by developmental regulator-like sequences (e.g., Sox14, Pnt, and GATAd-like motifs).

### Core Circadian Clock Genes Show Limited Transcriptional Rhythmicity

Canonical insect clock gene homologs showed limited transcriptional rhythmicity across all three detection approaches (Table S11). Under the primary model framework, rhythmic clock transcripts were detected only in wheat-fed aphids, and only four transcripts were assigned to a rhythmic class, mostly under LD: specifically, timeless X1 (dark-induced), and cycle X3, shaggy X4, and casein kinase IIα X2 (all diurnally rhythmic, WhR2). No clock isoforms were found to be rhythmic on artificial diet, consistent with the broader loss of rhythmicity in the absence of host cues. Cosinor and JTK_CYCLE identified additional, nominally rhythmic clock isoforms under specific conditions, particularly on wheat, but these signals were sparse and inconsistent across methods.

## Discussion

In this study, we utilized a 48 h time course to characterize the diurnal and circadian transcriptome of *R. padi* under four conditions spanning two light regimes (LD vs. DD) and two food sources (wheat vs. artificial diet). Across all tools, only a small fraction of the transcriptome exhibited 24 h rhythms, and this fraction notably weakened as zeitgebers were removed, from 4.6% on wheat to 1.6% on diet as detected by dryR. Unexpectedly, rhythmicity was higher in DD than under LD conditions on wheat, a pattern absent on artificial diet. Further, rhythmic transcripts on both hosts showed a consistent bimodal phase distribution near dawn and pre-dusk, with amplitude declining as external cues were removed. Functional enrichment linked rhythmic transcripts broadly to pathways related to RNA processing and metabolism. Promoter analysis found no strong E-box enrichment among rhythmic transcripts but identified recurring Mad/SMAD-, bZIP-, and zinc-finger-like motifs. The canonical clock gene homologs themselves showed limited 24 h transcriptional rhythmicity, detected only in wheat under LD conditions. In a complementary performance assay, survival was unaffected by host or light treatment. At the same time, reproductive output was reduced in wheat fed aphids reared in DD and in those reared on artificial diet. While the exact number and identities of rhythmic transcripts varied across statistical methods and significance thresholds, these shared trends indicated limited, cue-dependent transcriptional rhythmicity in *R. padi* at the whole-body level.

Two hypotheses may explain our results and are not mutually exclusive. First, the decline in both the prevalence and amplitude of rhythmic transcripts as external cues were removed is well explained by a damped-oscillator model proposed for the aphid circadian clock, in which endogenous rhythms decrease in amplitude, and in some cases lost altogether, under free-running conditions [3, 16, 21, 55]. Second, sampling whole-body rhythmicity may additionally limit our ability to detect rhythmicity that may be spatially restricted to particular tissues or cell types. For example, canonical clock genes are typically transcribed in a small population of specialized neurons rather than throughout the entire brain or body, and transcriptional rhythms related to metabolic processes have been shown to differ between tissues [18–21, 56–58]. By sampling at the whole-body level, expression is averaged across heterogeneous tissues, effectively diluting and potentially destructively interfering with any localized rhythmic signal. This may explain why the proportion of transcriptional rhythmicity for core clock genes was rather low. Further, the transcripts detected as rhythmic in this analysis are likely biased toward genes that are broadly and homogeneously transcribed across the body, as these would be the signals most likely to remain detectable after summing expression across the whole body.

Initially, we hypothesized that overall transcriptional rhythmicity would decrease in DD conditions, independent of host treatment, as light is thought to be the primary zeitgeber for most insects [59]. However, dryR and Cosinor (at the *p* < 0.05 level) revealed a higher level of rhythmic transcripts when aphids fed in DD compared to LD on wheat, a trend that was not seen in diet-fed aphids. The absence of heightened DD rhythmicity in diet-fed aphids implicates the host plant, rather than the aphids’ endogenous clock, as the primary driver of this rhythmicity. One explanation is that rhythmic changes in phloem quality [60, 61], driven by the plant’s clock continuing to oscillate under DD conditions, entrain a subset of transcriptional rhythms in the aphid which not typically rhythmic under LD. Consistent with this, dark-induced rhythmic transcripts on wheat displayed an approximately 4 h phase delay relative to diurnally rhythmic transcripts on wheat. This delay was absent from dark-induced transcripts from diet-fed aphids, which retained phase distributions like those observed in LD, although it should be noted that the number of dark-induced transcripts on diet was comparatively small (*n* = 10). However, if the aphid’s free-running clock were responsible for the phase shift, we would expect to see a comparable phase delay for diet-fed aphids in DD conditions, which was not observed.

Further, functional enrichment in dark-induced transcripts on wheat highlights protein folding, amino acid metabolism, and carbon metabolism, categories which are consistent with an aphid transcriptional response to metabolic shifts in host physiology that accompany sustained darkness, including reduced photosynthetic output and carbon transport and partitioning [62–66]. Together, these patterns suggest that rhythmic plant-derived stimuli likely persist under DD conditions and continue to shape aphid transcriptional rhythmicity independently of light-dark cycles. However, this heightened DD-rhythmicity on wheat appears fragile, as it was not supported by JTK_CYCLE or Cosinor after FDR correction (*q* < 0.05). This signal being captured only by dryR and Cosinor may suggest that dark-induced transcripts on wheat exhibit a weak or noisy oscillatory structure that is preferentially detected by harmonic regression-based methods rather than the non-parametric approach of JTK_CYCLE (Supplemental Note S1).

The transcriptional signatures of metabolic stress in DD on wheat are further supported by the performance assay (Fig. S3). While adult survival did not significantly differ across any of the four treatment combinations over the 96 h window, reproductive output did. Aphids reared in LD on wheat produced substantially more offspring than any other treatment, while aphids reared in DD on wheat produced significantly fewer nymphs than diet-fed aphids in DD, though not significantly fewer than diet-fed aphids in LD. In contrast, aphids on an artificial diet showed no significant difference in either survival or reproduction between LD and DD. Because both treatments shared nearly identical regimes, including a constant temperature in DD versus a 25/19 °C diurnal temperature cycle in LD, the reproductive penalty seen on wheat indicates that darkness, and the resulting lack of temperature cycles, act primarily by degrading host plant quality rather than directly stressing the aphid. Furthermore, diet-fed aphids sustained comparable performance. Additionally, components in artificial diets are known to degrade in direct light, which may explain the trend toward higher reproduction in DD. Reduced transcriptional rhythmicity on diet is therefore unlikely to reflect poor diet quality or generalized aphid stress, but rather the absence of rhythmic, plant-derived cues.

Beyond the total number of rhythmic transcripts, the temporal organization and amplitudes of these rhythms across light and host-treatments further point to the strong cue-dependency of the aphid clock. We observed a consistent bimodal phase distribution across most rhythmicity models assigned by dryR, with peak transcript expression clustering around dawn (ZT/CT0) and pre-dusk (ZT/CT12). These peaks likely represent critical periods when aphids synchronize their physiology with predicted environmental conditions or host-plant metabolic shifts that are ecologically relevant for coordinating feeding, detoxification, and resource allocation. Further, we detected a clear decline in rhythmic amplitude as cues were removed, with the lowest amplitudes observed in constant darkness on artificial diet. This concurrent decrease in both the abundance and the magnitude of oscillations across host and light treatments is consistent with a damped-oscillator model, though formally distinguishing damping from simple cue dependence would require tracking amplitude across multiple consecutive DD cycles and likely a higher sampling resolution.

The specific transcripts identified as rhythmic in this study provide a foundation for understanding the role that rhythmic transcription plays in aphid biology. Components of the canonical insect clock, including cycle, shaggy, and casein kinase IIα, displayed diurnal rhythms when aphids fed on wheat. Most other known insect clock genes failed to remain rhythmic after filtering or FDR correction thresholds in any light or host condition. Only a few known clock genes exhibiting rhythmicity exclusively under light-dark cycles on wheat provide further evidence of a weak oscillator that relies heavily on environmental reinforcement. In addition, these genes are often expressed rhythmically in specialized neuronal clusters in the brain, making rhythmic detection challenging in whole-body samples as discussed previously. This underscores the need for tissue-level datasets to better resolve rhythmicity in key regions such as the brain, salivary gland, and gut [18–21, 56–58]. The tissue-dilution effect is not limited to clock genes; it applies broadly to any transcript whose rhythmicity is restricted to specific cells and tissues. The modest whole-body rhythmicity reported here should therefore be interpreted as a lower bound on the actual extent of circadian transcriptional regulation in *R. padi*.

Among the transcripts that maintained rhythmicity across both host treatments, polycomb-like (XM_060995664.1) and neprilysin-2 (XM_060991018.1) emerge as particularly strong candidates for future study. Polycomb-like’s shared circadian rhythmicity across both wheat and artificial diet is consistent with its established role in regulating sleep-wake cycling in Drosophila [49, 50], while neprilysin-2’s circadian rhythmicity across both hosts, despite a shifted phase between LD and DD on wheat, aligns with its implicated roles in feeding behavior and clock output in other insects [51, 52]. The persistence of these transcripts under free-running conditions on both wheat and diet suggests they are under direct circadian control and may link the aphid clock to core physiological and behavioral processes such as feeding, metabolism, locomotion, and reproduction. Disrupting these clock-linked pathways, especially those with putative roles in the plant-aphid relationship, through targeted approaches like RNAi, could provide a novel strategy for aphid control [4, 5].

Beyond individual candidate genes, functional enrichment analysis revealed a large number of rhythmic transcripts involved in RNA metabolism, transcription initiation, and mRNA splicing, particularly under LD conditions. The rhythmicity of these pathways suggests that a significant part of the aphid’s temporal organization is dedicated to regulating RNA and transcription, which is consistent with a growing body of work showing that pre- and post-transcriptional regulation, including alternative splicing and RNA stability, is an important part of the circadian regulatory system [67]. Our results suggest that aphids may utilize similar rhythmic transcriptional and splicing machinery to coordinate physiological states across the whole body.

To identify the regulatory mechanisms of *R. padi* rhythms, we examined promoter regions for both canonical and novel motifs. Interestingly, E-box motifs (CACGTG/CANNTG) were found in approximately 10% of both rhythmic and non-rhythmic transcripts, suggesting that an E-box alone is likely insufficient to drive rhythmic expression. However, the subset of rhythmic transcripts (∼10–17% depending on rhythmic class) that carry E-box motifs remain candidates for direct CLK/CYC regulation, representing a putative tier of direct clock targets embedded within a larger set of indirectly regulated outputs. Further, the lack of consistent E-box enrichment across rhythmic transcripts suggests that whole-body transcriptional rhythmicity in *R. padi* may be primarily shaped by regulatory mechanisms downstream of the core clock’s canonical E-box-binding transcription factors.

Our *de novo* promoter motif discovery identified several recurring signatures, including significantly enriched GC-rich (Mad/SMAD-like), bZIP, and C2H2 zinc finger (Krüppel-type) motifs, each of which has been implicated in circadian regulation [68–72]. Because clock-controlled gene expression in insects is often mediated indirectly, through transcription factors and regulatory elements downstream of the core CLK/CYC E-box binding, the enriched Mad, zinc-finger, and bZIP motifs identified here may be a part of this downstream regulatory mechanism. We also detected enrichment of hairy-like bHLH motifs, which is notable because Hairy/HES-related bHLH-orange factors can recognize E-box or E-box-like sequences. In insects, the best-supported circadian member of this broader family is clockwork orange, an E-box-binding repressor that modulates clock transcription in Drosophila and other insects [73–75]. Thus, these hairy-like motifs may reflect variants of E-box regulatory domains in aphids that differ slightly from those in model insects.

In summary, our results indicate that whole-body transcriptional rhythmicity in *R. padi* is modest and strongly cue-dependent, consistent with a damped oscillator that relies on external, and possibly largely host-derived, reinforcement rather than a robust, self-sustaining one. Several limitations complicate this interpretation. Because rhythmicity was assessed in whole bodies, our estimates likely represent a lower bound on tissue-level circadian regulation, particularly for canonical clock genes expressed in restricted neuronal populations. Further, because the wheat and diet experiments were conducted on different dates and could not be directly statistically compared, differences between host treatments should be interpreted as descriptive rather than formally tested. Additionally, the transition period between light removal and the onset of free-running rhythmicity has not been sufficiently characterized in *R. padi* or aphids in general. As such, we cannot exclude the possibility that our 48 h DD entrainment window captured this transition period rather than a true free-running state. We chose this 48 h duration because it was the longest practical window given the experimental constraints. Any longer exposure to constant darkness for wheat-fed aphids risks greater physiological stress and possible impacts on adult survival. Further, a longer duration would have caused the artificial diet, which we did not replenish during our entrainment and sampling windows to limit disruption, to continue to diminish in quality and risk becoming contaminated.

## Conclusions

This study indicates that whole-body transcriptional rhythmicity in *R. padi* is limited in scope and appears strongly dependent on external cues, particularly those that may be derived from the host plant, rather than reflecting a robust, self-sustaining endogenous oscillator. Rhythmicity declined as light and host cues were removed, was organized around a consistent dawn and pre-dusk phase structure, and was most pronounced among transcripts linked to RNA processing and metabolism, and specifically among wheat-fed aphids in constant darkness, to protein folding and carbon/amino-acid metabolism. This latter signature, together with the reduced reproductive output observed specifically in wheat-fed aphids reared in DD, is consistent with a host-plant nutritional-stress cascade linking the absence of light for photosynthesis to aphid performance, though this hypothesis requires direct testing. Promoter analyses further identified candidate regulatory motifs, including Mad/SMAD-, bZIP-, and zinc-finger-like elements, that may act downstream of the core clock’s E-box-binding machinery, and a small set of transcripts, including polycomb-like and neprilysin-2 orthologs, emerged as strong candidates for direct circadian regulation across both host treatments.

Together, these findings position *R. padi* as a system in which whole-body transcriptional rhythms are shaped more by the daily and metabolic state of the host plant than by a strong endogenous clock, while identifying specific transcripts, pathways, and regulatory motifs for future mechanistic study. Because whole-body sampling likely underestimates rhythmicity concentrated in specific tissues, tissue-resolved transcriptomics of the brain, gut, and salivary glands represents the most direct next step for testing whether a stronger endogenous oscillator is present but diluted in our data. Further, RNAi-mediated knockdown of high-priority candidates such as polycomb-like and neprilysin-2, together with high-throughput behavioral tracking, could help establish whether these rhythms causally contribute to aphid feeding, reproduction, and host colonization and whether any rhythms exist in these aspects of aphid life. Given the global distribution and economic impact of *R. padi* as a cereal pest and virus vector, resolving how host and light cues shape its daily biology may ultimately help inform the timing and improve pest management interventions.

## List of Abbreviations

ANOVA: analysis of variance
BICW: Bayesian inference criterion weight
bHLH: basic helix-loop-helix
bZIP: basic leucine zipper
C2H2: Cys2-His2 zinc finger
CLK/CYC: CLOCK/CYCLE
CSU: Colorado State University
CT: circadian time
DD: constant darkness
DNase: deoxyribonuclease
FDR: false discovery rate
GC: guanine-cytosine (content)
GO: Gene Ontology
HSD: honestly significant difference
KEGG: Kyoto Encyclopedia of Genes and Genomes
LD: light-dark
mRNA: messenger RNA
NCBI: National Center for Biotechnology Information
NSF: National Science Foundation
PDF: pigment-dispersing factor
Pnt: Transcription factor Pointed
RNAi: RNA interference
RNA-seq: RNA sequencing
SRA: Sequence Read Archive
TSS: transcription start site
VST: variance-stabilizing transformation
ZT: zeitgeber time
WhR0–WhR5, AdR0–AdR5: dryR rhythmicity models for wheat (Wh) and artificial diet (Ad), respectively. R0 = excluded for low model support; R1 = arrhythmic; R2 = diurnal-only; R3 = dark-induced; R4 = shared-rhythm (uniform across LD and DD); R5 = differentially rhythmic (phase and/or amplitude differs between LD and DD)
WhM1/WhM2, AdM1/AdM2: dryR mean models. M1 = shared midpoint expression between LD and DD; M2 = differential midpoint expression

## Declarations

### Ethics approval and consent to participate

Not applicable.

### Consent for publication

Not applicable.

### Availability of data and materials

Raw RNA-seq reads and associated metadata are publicly available in the NCBI Sequence Read Archive (SRA) under BioProject accession PRJNA1413204. Code for RNA-seq processing and transcript quantification can be accessed at: https://github.com/dankunk/tango-public/. R code for downstream analyses, including dryR modeling and functional enrichment, is available at: https://github.com/dankunk/Rpadi_Wheat-Diet_LD-DD_Analysis, while promoter analysis scripts are provided at: https://github.com/dankunk/RhythmicPromoterAnalysis. All supplemental material can be found at: https://zenodo.org/records/23250260.

### Competing interests

The authors declare no potential conflicts of interest with respect to the research, authorship, and/or publication of this article.

### Funding

This work was supported by the U.S. Department of Agriculture, National Institute of Food and Agriculture Predoctoral Fellowship [award no. 2024-67011-42995] and start-up funds awarded to Vamsi Nalam from CSU.

### Authors’ contributions

D.K. and V.N. conceived and designed the study. D.K. conducted all experiments, performed bioinformatic analyses, drafted the manuscript, and contributed to funding acquisition. V.N. provided supervision, resources, and funding acquisition. Both authors interpreted results and revised the manuscript.

## Acknowledgements

The authors thank Josie Weese (Colorado State University - CSU) for her help with plant growth and colony maintenance. This work utilized the Alpine high-performance computing resource at the University of Colorado Boulder, funded by the University of Colorado Boulder, the University of Colorado Anschutz, and Colorado State University, with support from NSF grants OAC-2201538 and OAC-2322260. The authors used ChatGPT-5 (OpenAI), Grammarly, and Claude Sonnet-5 (Anthropic) to assist with language editing and manuscript preparation. The authors take full responsibility for the content of the manuscript.

## Supplemental Figures and Legends

**Fig. S1.**
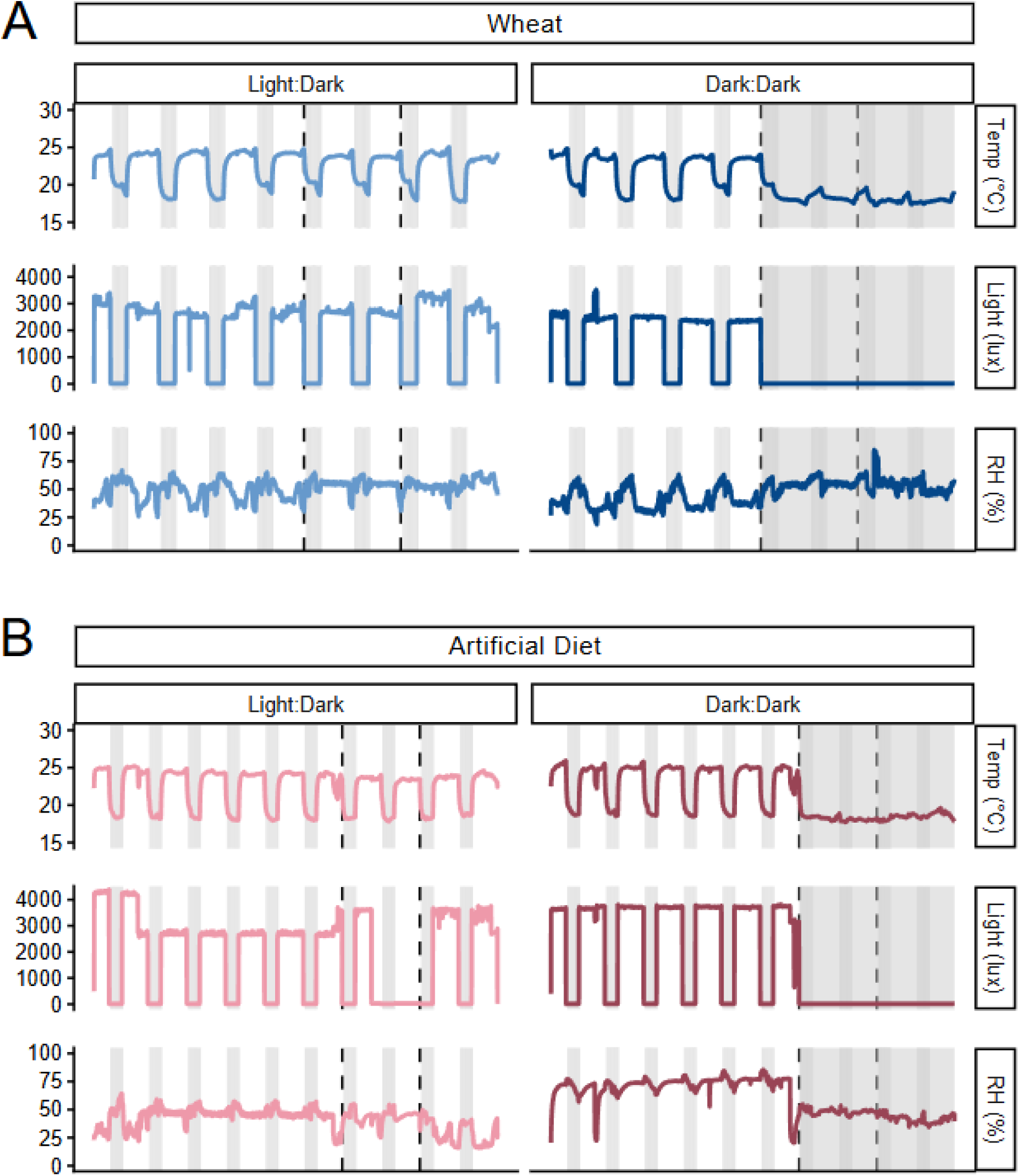
Environmental conditions during entrainment and sampling. Environmental conditions recorded during wheat (A) and artificial diet (B) experiments under LD (left) and DD (right). Lines show temperature (°C), light intensity (lux), and relative humidity (RH%). Background shading indicates the light schedule (white = photophase; gray = scotophase). In DD panels, additional shading indicates the transition to constant conditions and the corresponding subjective day and night. Dashed vertical lines mark the start and end of the 48 h entrainment window. In the LD condition of the artificial diet, the environmental monitor failed to record light intensity during part of the entrainment window. Personnel monitoring confirmed that the lights were operating normally during this period, consistent with the diurnal cycling of temperature and the maintenance of daytime temperatures. Missing light values were not interpolated; instead, the gap is presented as recorded.

**Fig. S2.**
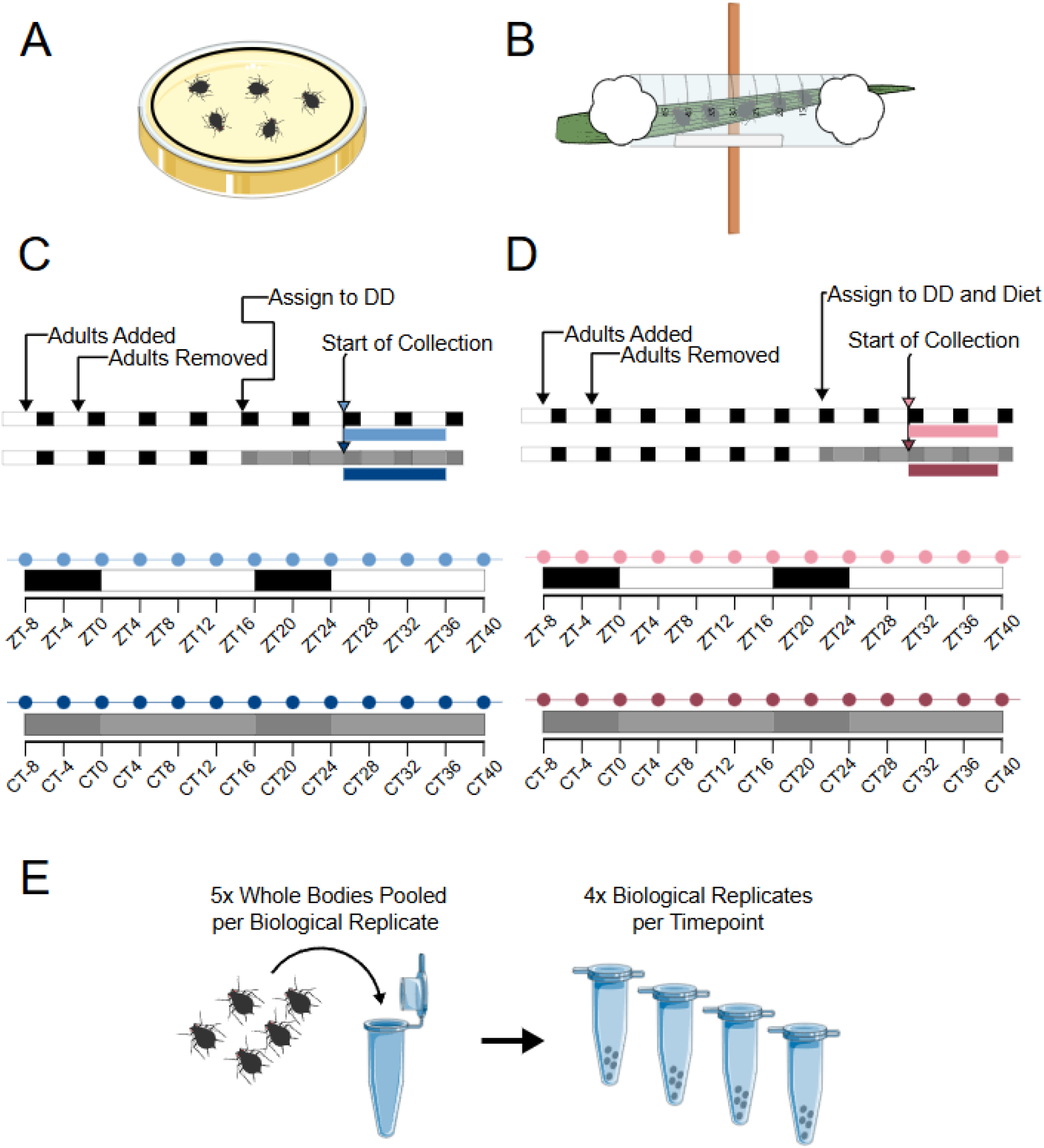
Experimental design and sampling scheme. Overview of experimental setup and sampling. Diagram of the (A) artificial diet sachets and (B) wheat chambers utilized in each experiment. (C, D) Wheat and artificial diet experimental timelines showing age-synchronization, transfer to DD, and the start of sampling (arrows). White and black bars indicate the light schedule (white = photophase; black = scotophase). Gray shading indicates subjective day/night. Light blue/red points indicate sampling times in LD (zeitgeber time, ZT) and dark blue/red points indicate DD (circadian time, CT). (E) RNA-seq sampling strategy: five whole bodies were pooled per biological replicate, with four biological replicates collected at each timepoint.

**Fig. S3.**
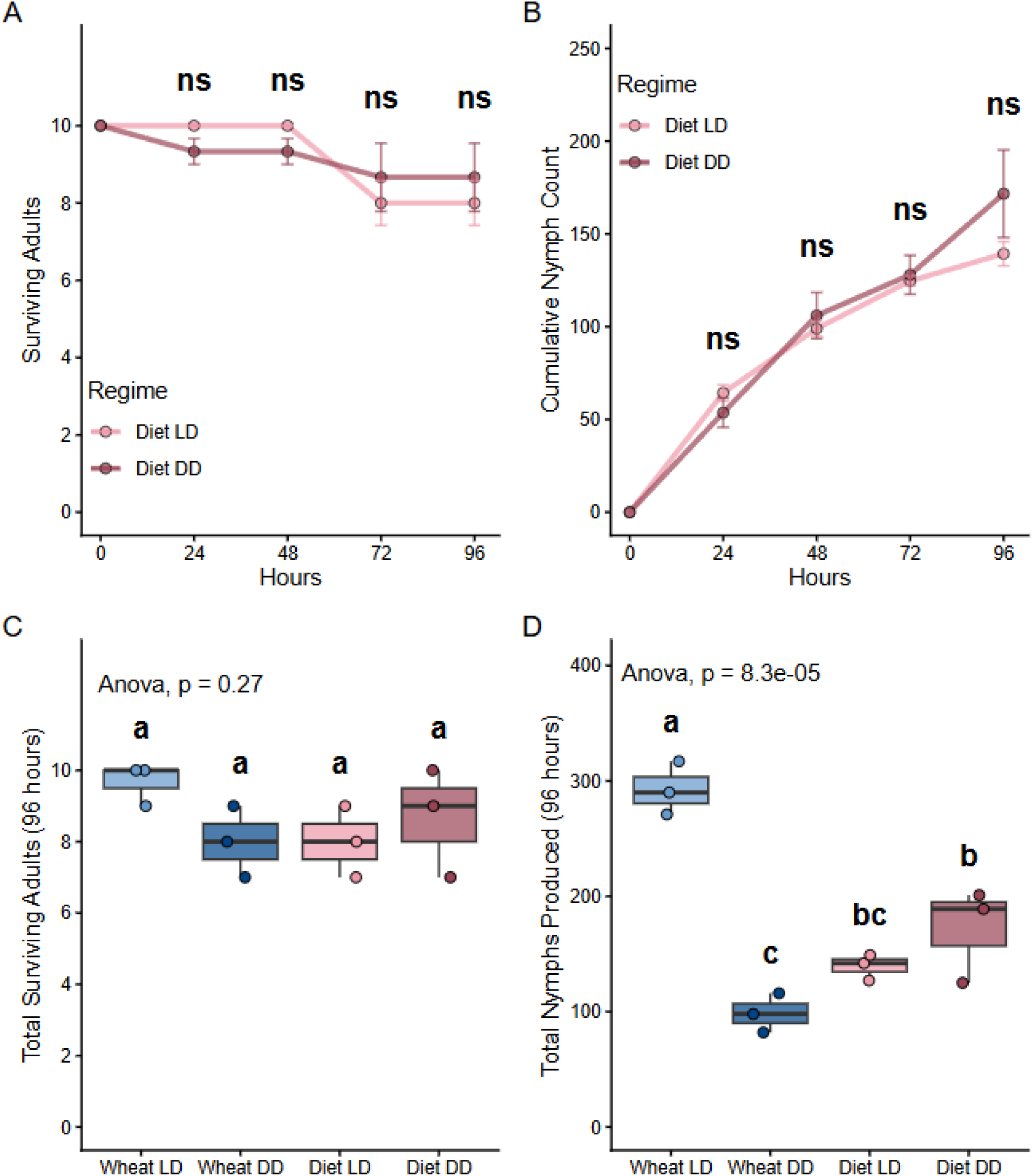
Aphid survival and reproductive performance across host and light regimes. Survival and reproductive performance assays of *R. padi* across 96 h in all host and light treatments. (A) Daily number of surviving adults and (B) cumulative number of nymphs produced by aphids feeding on artificial diet under LD and DD; “ns” indicates no significant differences between light regimes at a given timepoint (t-test). (C) Total surviving adults and (D) total nymphs produced across all four treatment combinations at the 96 h timepoint. Boxplots show the median and interquartile range with individual cage and sachet replicates displayed as jitter points. Letters denote statistically significant group differences (Tukey’s HSD, *α* = 0.05) following a significant one-way ANOVA, with each corresponding p-value annotated on each panel.

**Fig. S4.**
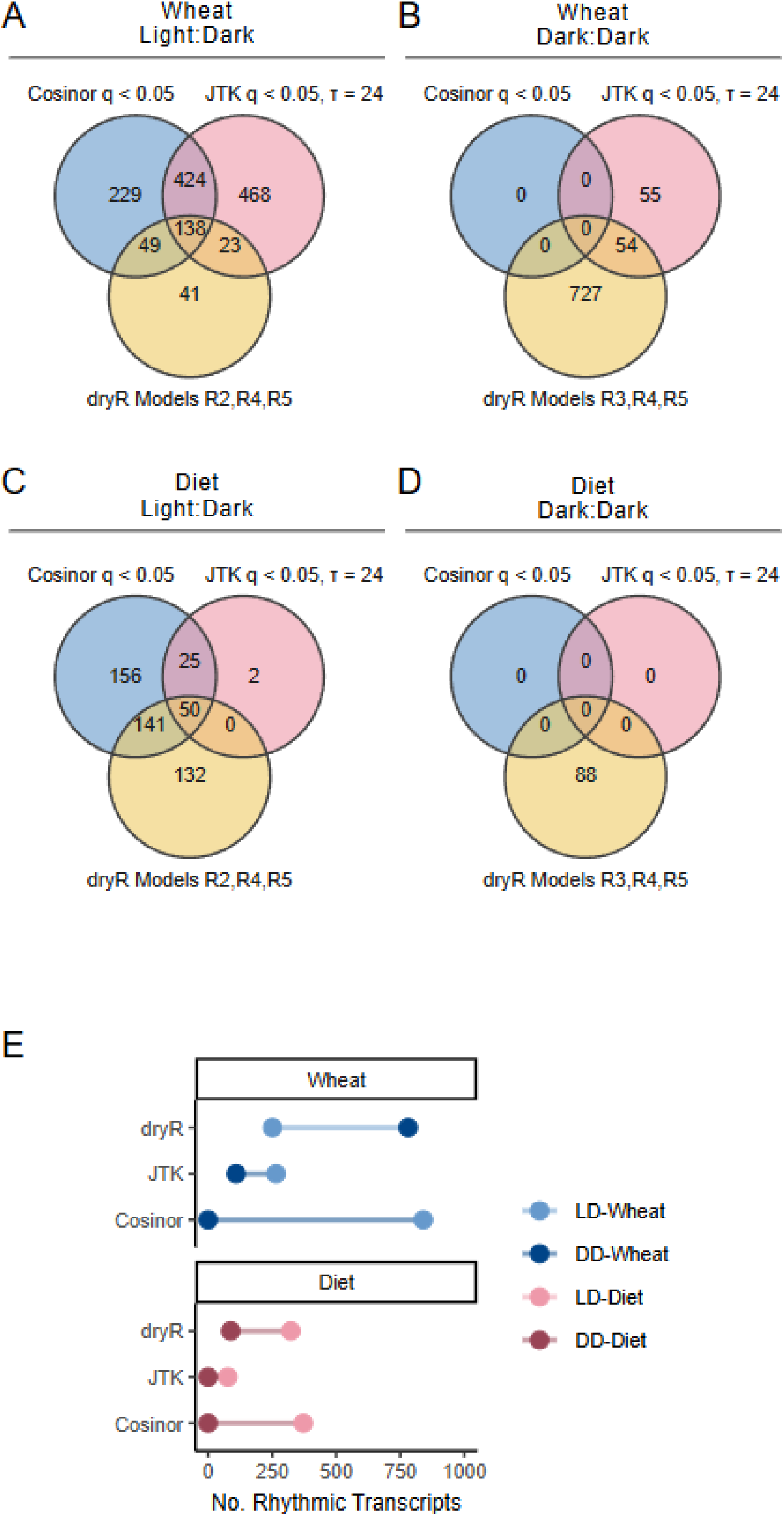
Comparison of rhythmic transcript detection across methods. Venn diagrams showing overlap among rhythmic transcripts identified by Cosinor (*q* < 0.05), JTK_CYCLE (*τ* = 24 h, *q* < 0.05), and dryR under LD and DD conditions in wheat (A-B) and artificial diet (C-D). For dryR, rhythmic transcripts in LD include models R2, R4, and R5, while in DD, they include models R3, R4, and R5. Numbers indicate transcript counts for each method and their corresponding intersections. (E) Dumbbell plots showing the number of rhythmic transcripts detected under LD versus DD for all three rhythmicity methods (Cosinor < 0.05, JTK_CYCLE < 0.05, and dryR R2-R5) in the wheat (top) and artificial diet (bottom) experiments. Dots indicate the number of rhythmic transcripts in LD (light shades) and DD (dark shades), and connecting lines show the difference between counts in LD and DD for each method within each experiment.

**Fig. S5.**
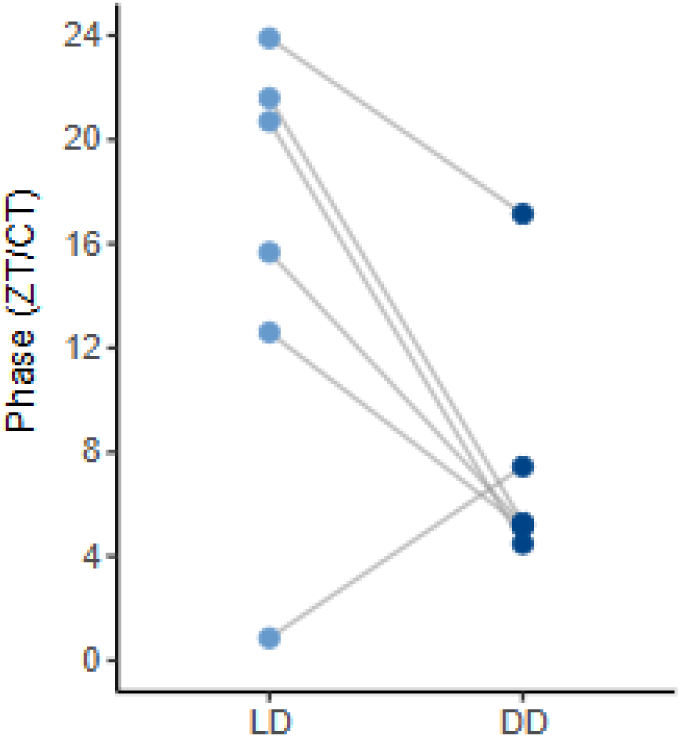
Rhythmic model R5 phases in wheat under LD and DD. Paired Dumbbell plot showing phase under LD (ZT) and DD (CT) conditions for transcripts assigned to dryR model R5 in the wheat experiment. Each dot represents the transcript phase, and connecting lines illustrate the phase difference between LD and DD.

**Fig. S6.**
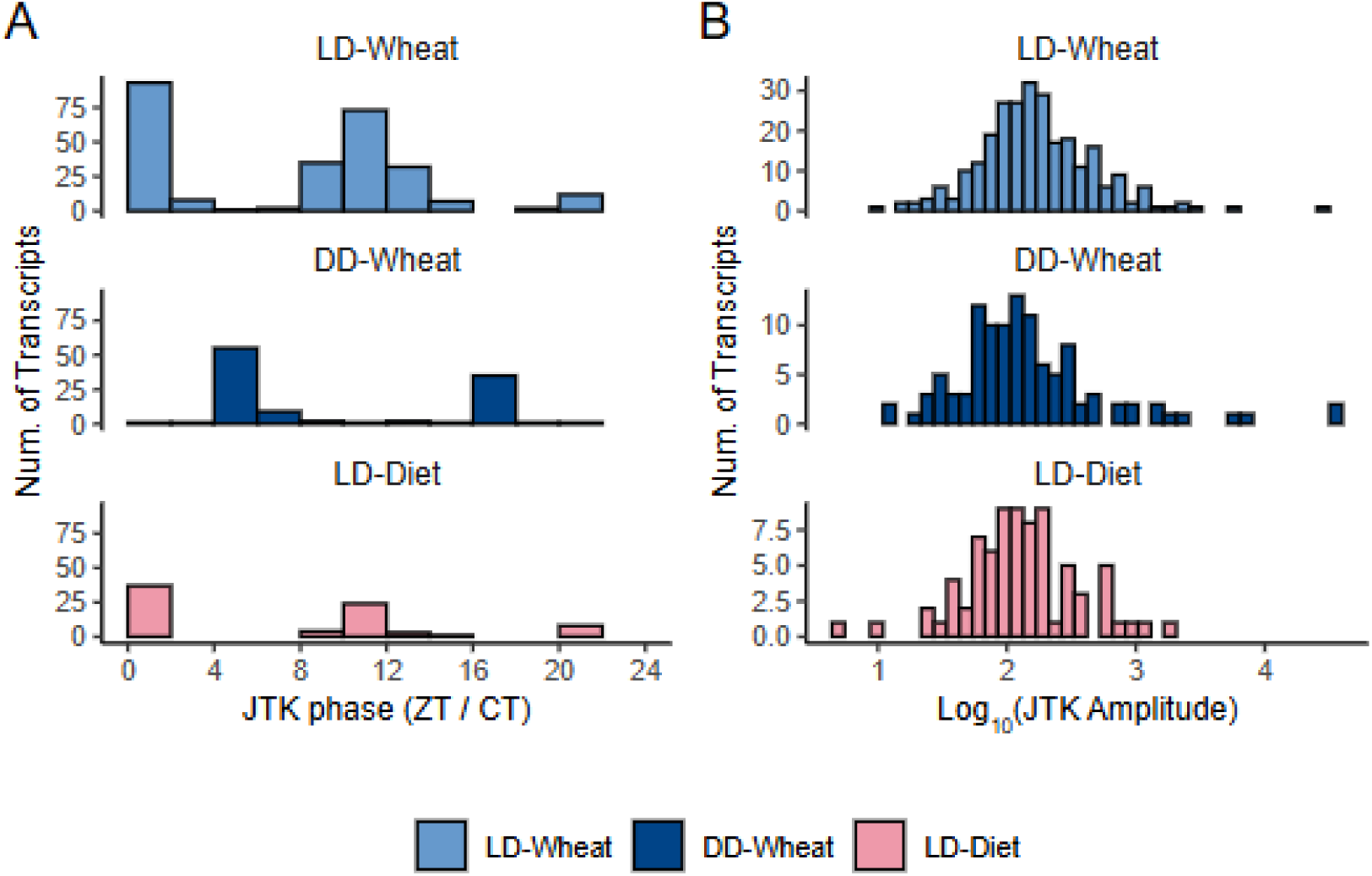
JTK_CYCLE phase and amplitude distributions. Histograms showing JTK_CYCLE rhythmic parameters for transcripts with JTK *q* < 0.05 and τ = 24 h, for each experimental condition. (A) Phase distributions for transcripts rhythmic in LD (ZT) and DD (CT). (B) Amplitude distributions are shown on a log_10_-transformed scale where amplitude is reported as the median sign-adjusted deviation. Colors indicate condition and experiment (LD–Wheat, DD–Wheat, LD–Diet).

**Fig. S7.**
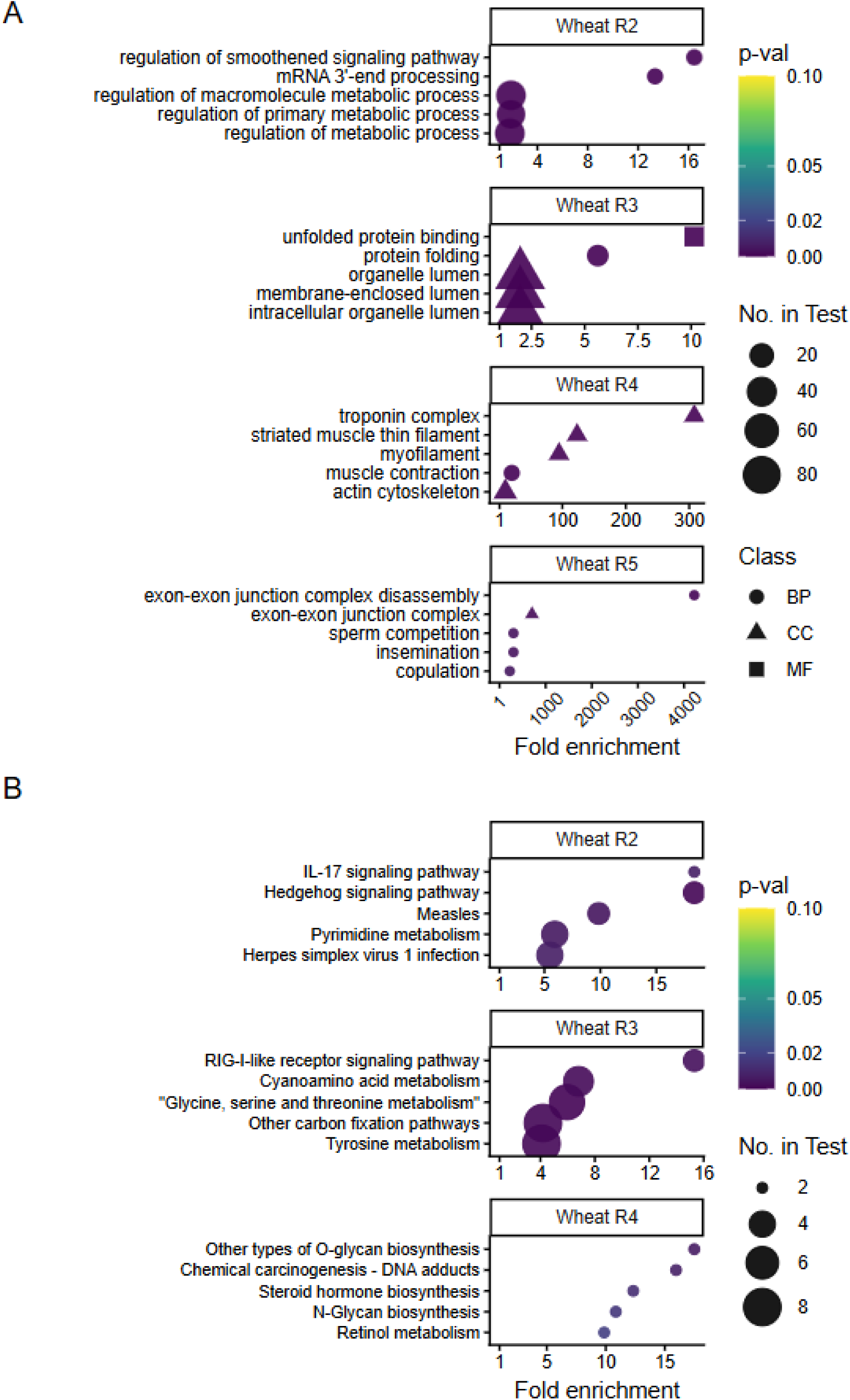
GO and KEGG enrichment of rhythmic transcripts across dryR rhythmicity models in wheat. Bubble plots showing (A) enriched Gene Ontology (GO) terms and (B) KEGG pathways for rhythmic transcripts in wheat across dryR rhythmicity models (R2-R5). The size of each bubble represents the number of rhythmic transcripts annotated to the term/pathway in the test set. Bubble color indicates Fisher’s exact test *p*-value. In GO plots, shape denotes GO domain: biological process (BP), cellular component (CC), or molecular function (MF).

**Fig. S8.**
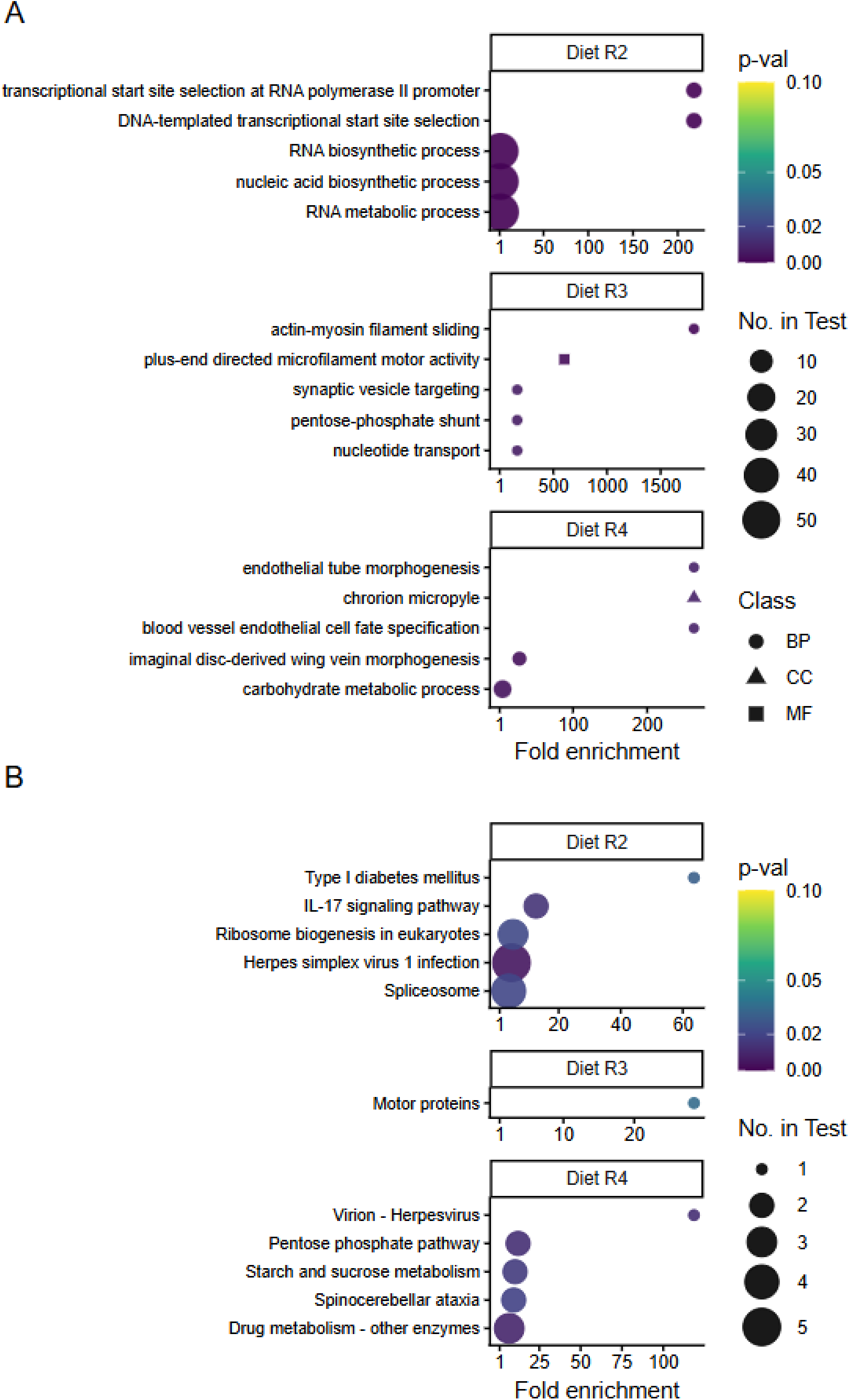
GO and KEGG enrichment of rhythmic transcripts across dryR rhythmicity models in artificial diet. Bubble plots showing (A) enriched Gene Ontology (GO) terms and (B) KEGG pathways for rhythmic transcripts on artificial diet across dryR rhythmicity models (R2 -R5). Bubble size represents the number of rhythmic transcripts annotated to the term/pathway in the test set. Bubble color indicates Fisher’s exact test *p*-value. In GO plots, shape denotes GO domain: biological process (BP), cellular component (CC), or molecular function (MF).

**Fig. S9.**
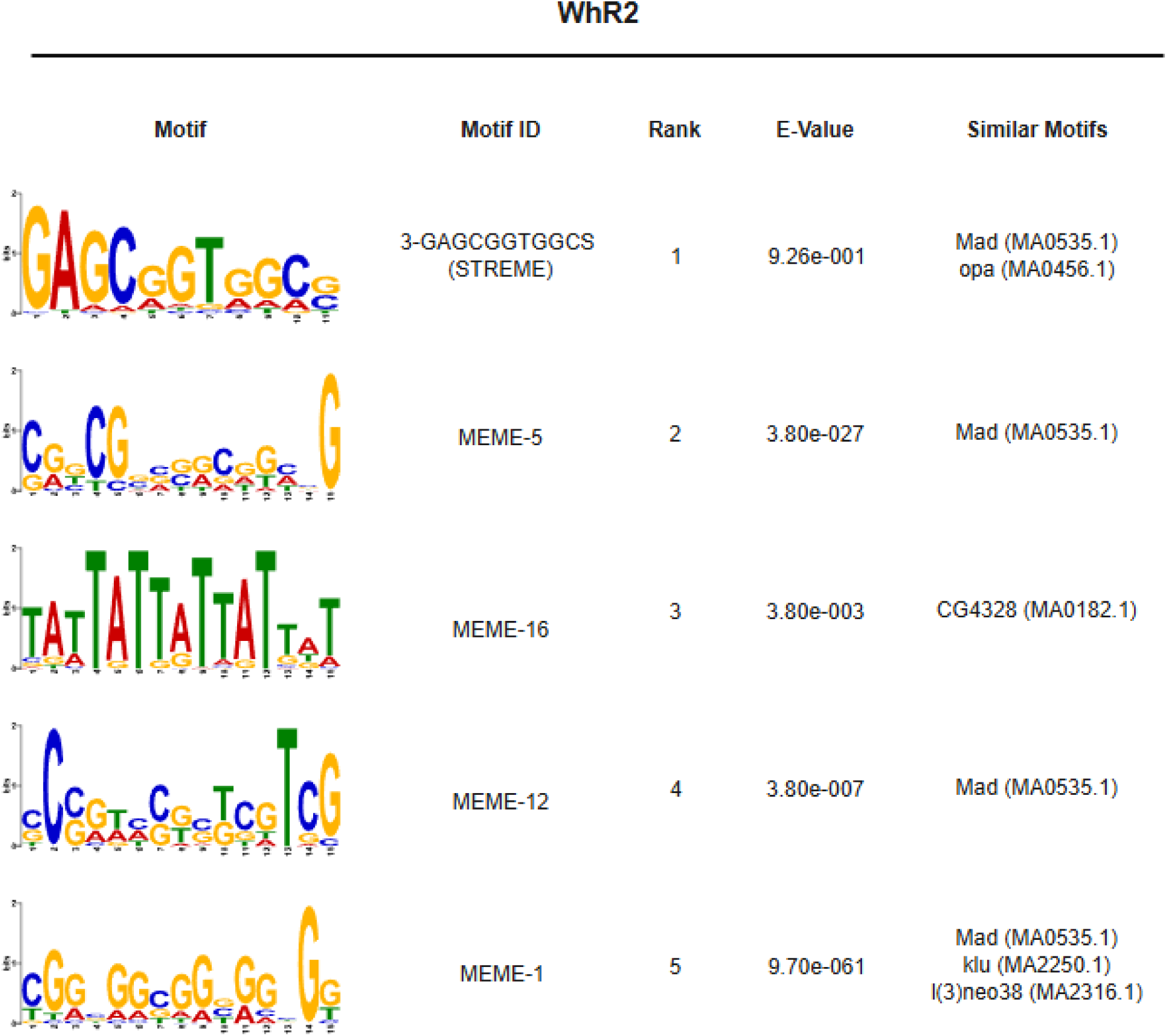
XSTREME motif enrichment among diurnal (WhR2) transcripts on wheat. Top five motifs identified by XSTREME among diurnal (WhR2) transcript promoters on wheat, relative to non-rhythmic (WhR1) promoters as background. Sequence logos represent position-specific nucleotide probabilities derived from MEME-format motifs, where letter height reflects nucleotide frequency at each position. Reported E-values correspond to XSTREME enrichment statistics; lower values indicate stronger enrichment. Complete XSTREME results, including SEA enrichment p-values and total motif occurrences, are provided in Table S10.

**Fig. S10.**
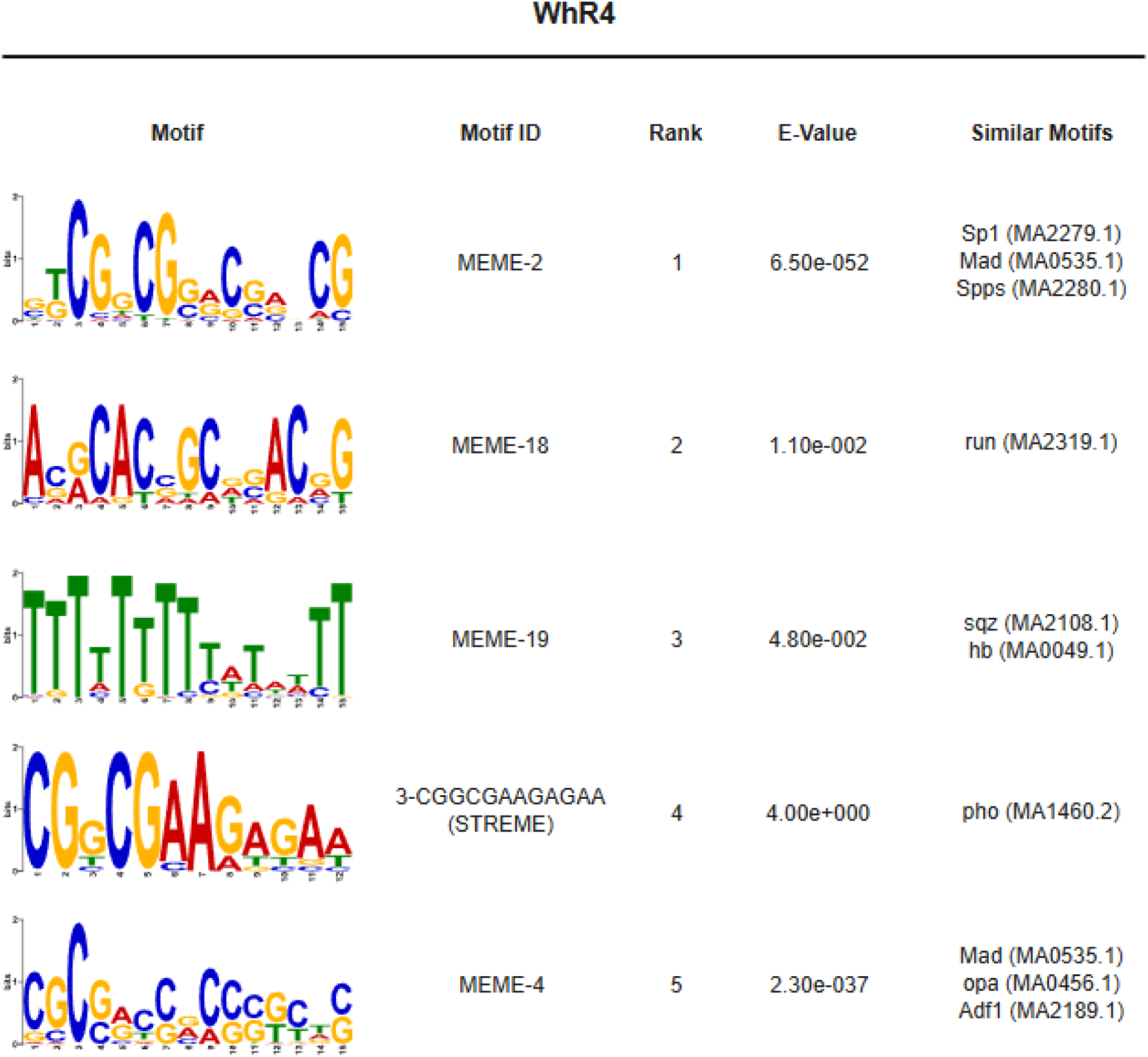
XSTREME motif enrichment among shared-rhythm (WhR4) transcripts on wheat. Top five motifs identified by XSTREME among shared-rhythm (WhR4) transcript promoters on wheat, relative to non-rhythmic (WhR1) promoters as background. Sequence logos represent position-specific nucleotide probabilities derived from MEME-format motifs, where letter height reflects nucleotide frequency at each position. Reported E-values correspond to XSTREME enrichment statistics; lower values indicate stronger enrichment.

**Fig. S11.**
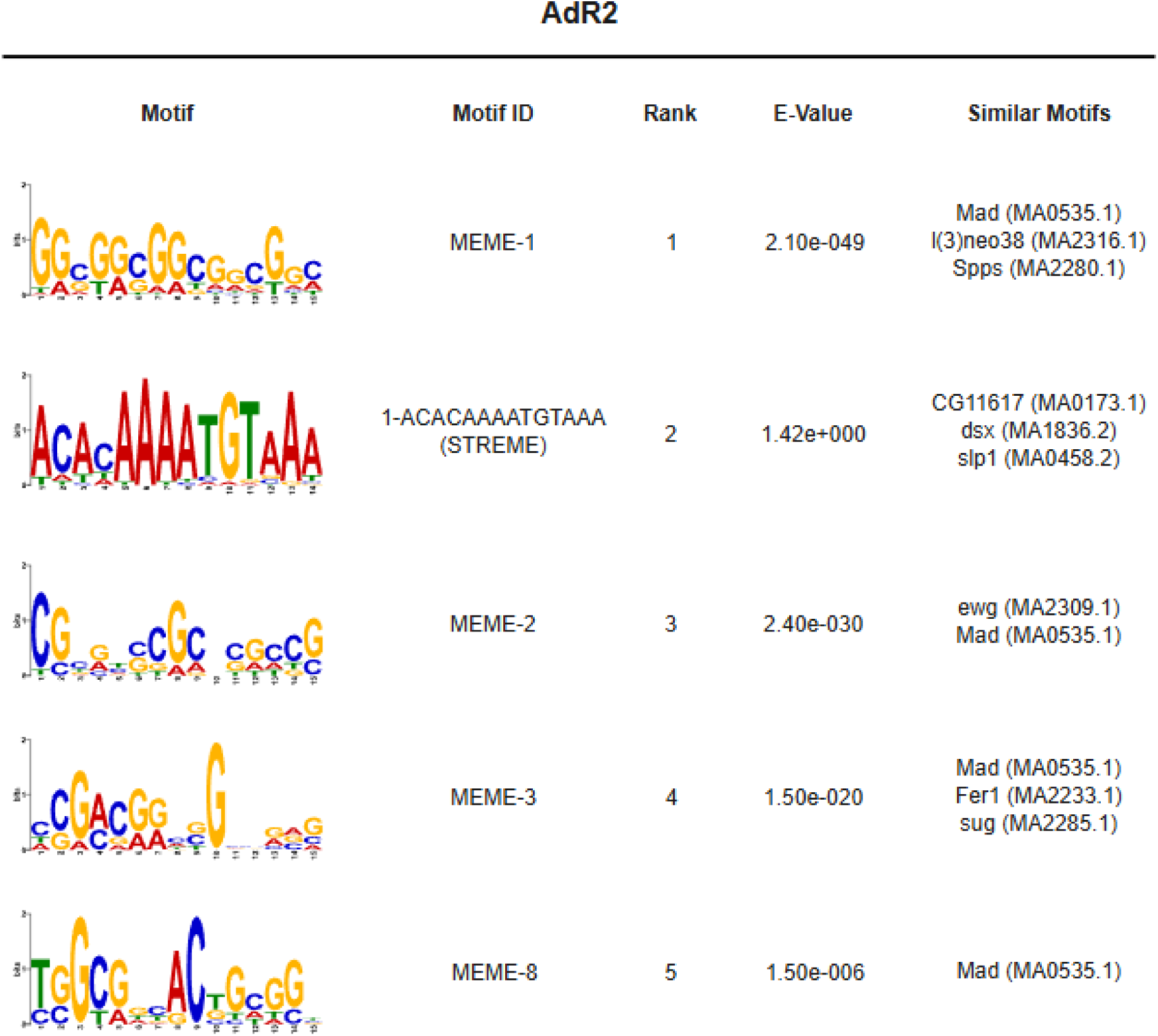
XSTREME motif enrichment among diurnal (AdR2) transcripts on artificial diet. Top five motifs identified by XSTREME among diurnal (AdR2) transcript promoters on artificial diet, relative to non-rhythmic (AdR1) promoters as background. Sequence logos represent position-specific nucleotide probabilities derived from MEME-format motifs, where letter height reflects nucleotide frequency at each position. Reported E-values correspond to XSTREME enrichment statistics; lower values indicate stronger enrichment.

**Fig. S12.**
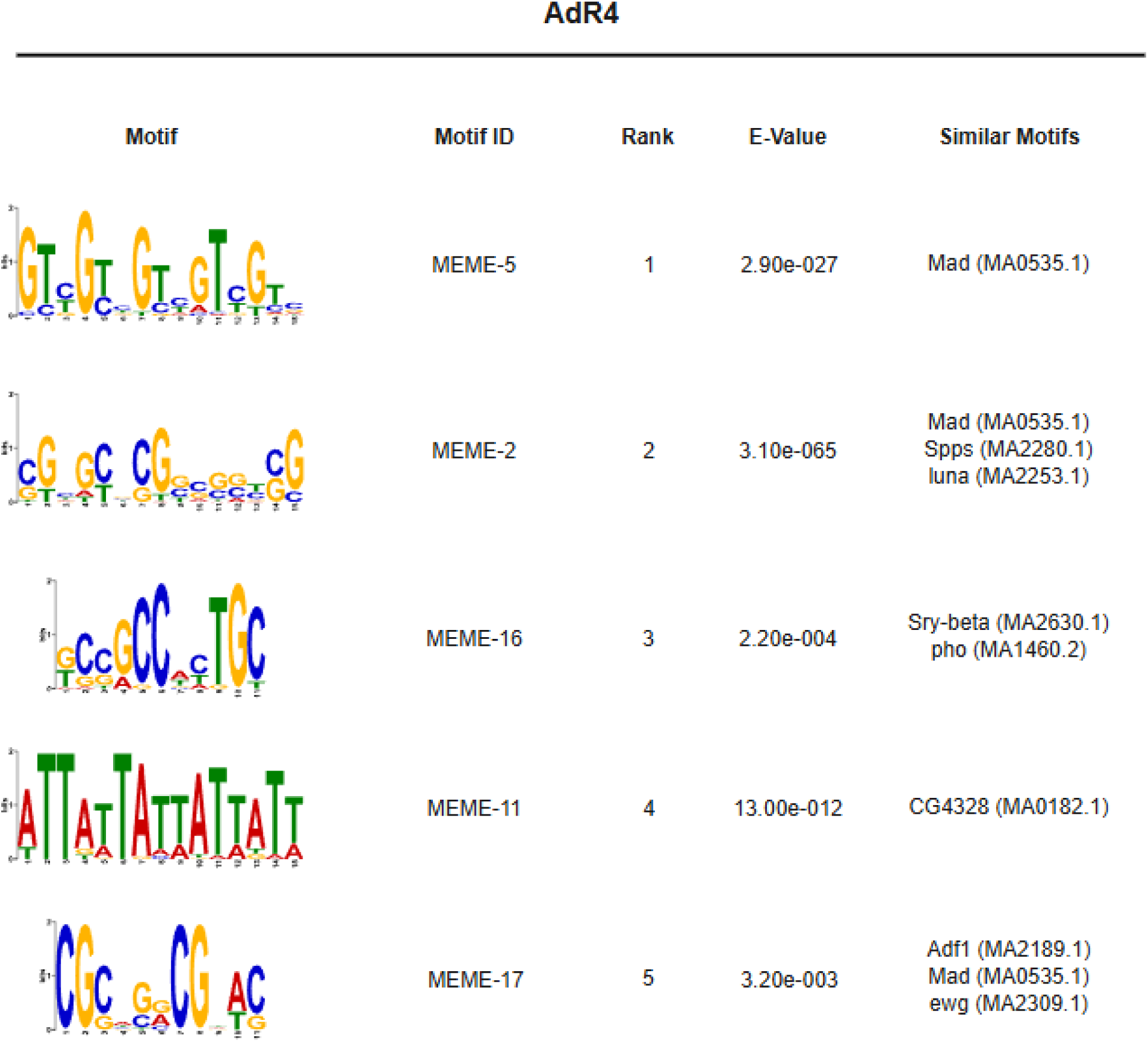
XSTREME motif enrichment among shared-rhythm (AdR4) transcripts on artificial diet. Top five motifs identified by XSTREME among shared-rhythm (AdR4) transcript promoters on artificial diet, relative to non-rhythmic (AdR1) promoters as background. Sequence logos represent position-specific nucleotide probabilities derived from MEME-format motifs, where letter height reflects nucleotide frequency at each position. Reported E-values correspond to XSTREME enrichment statistics; lower values indicate stronger enrichment.

## Supplemental Table Legends

**Table S1. Artificial diet composition and ingredients.**

Excel file containing the list of ingredients used to prepare the artificial diet, including supplier, catalog number, and amount added per 100 mL of diet. The table is organized by component class (sugar, amino acids, vitamins, and salts/other ingredients).

**Table S2. Survival and performance assay raw data and statistical results.**

Excel file containing the raw survival and performance assay datasets, including the daily counts of surviving adults and cumulative nymphs for diet-fed aphids and endpoint (96 h) counts for all treatments. Additional sheets report the full statistical results for each hypothesis test conducted.

**Table S3. RNA-seq quality metrics and sample metadata.**

Excel file containing per-sample MultiQC summary statistics for read preprocessing (fastp) and quantification (Salmon) (sheet 1) and sample metadata for each library (sheet 2), including FASTQ filenames, sample name, timepoint (ZT), biological replicate, sequencing batch, host treatment (i.e., wheat or diet), and light treatment (LD or DD).

**Table S4. dryR rhythmicity results, model summaries, and filtering statistics.**

Large Excel file containing dryR results for wheat and artificial diet experiments. dryR transcript-level results for wheat (sheet 1) and diet (sheet 2). Model-level summaries (sheet 3), including transcript counts per model and descriptive statistics for each experiment. Filtering summary for transcripts assigned to model R0 (sheet 4), including the reason(s) for filtering/low rhythmicity support.

**Table S5. Nitecap results.**

Large Excel file containing Nitecap results. Each sheet includes the input expression matrix and the Nitecap results for each experiment and light condition: LD – Wheat (sheet 1), DD – Wheat (sheet 2), LD – Diet (sheet 3), and DD – Diet (sheet 4).

**Table S6. dryR model comparisons and transition counts between wheat and artificial diet.**

Large Excel file summarizing dryR model correspondence for transcripts in wheat and artificial diet. This file contains the list of transcript model assignments between wheat and diet, excluding model R1 (sheet 1) and including R1 (sheet 2). Counts of model-to-model transitions from wheat to diet used in Sankey analysis (e.g., Wheat R1 → Diet R2) are also provided (sheet 3).

**Table S7. GO term enrichment results and associated transcript IDs for dryR rhythmicity models.**

Large Excel file containing Gene Ontology (GO) enrichment outputs for dryR rhythmicity models in wheat (R2–R5) and artificial diet (R2–R4). For each model, two sheets are provided: the first sheet contains GO enrichment results (starting with Wheat R2), and the following sheet lists the transcript IDs associated with each enriched GO term from the preceding sheet. In total, the workbook contains 14 sheets (two per model).

**Table S8. KEGG pathway enrichment results and associated transcript IDs for dryR rhythmicity models.**

Large Excel file containing KEGG pathway enrichment outputs for dryR rhythmicity models in wheat (R2–R5) and artificial diet (R2–R4). For each model, two sheets are provided: the first sheet contains KEGG enrichment results (starting with Wheat R2), and the following sheet lists the transcript IDs associated with each enriched pathway from the preceding results sheet. In total, the workbook contains 14 sheets (two per model).

**Table S9. FIMO motif scan outputs for dryR rhythmicity models.**

Excel file containing FIMO results from promoter motif scanning across dryR rhythmicity models in wheat (R1–R5) and artificial diet (R1–R4). Meme file motif definitions used for the scan are provided on sheet 1. Subsequent sheets contain the FIMO results tables for each model in wheat and diet.

**Table S10. XSTREME motif discovery results for dryR rhythmicity models.**

Excel file containing XSTREME output tables for promoter motif discovery across dryR rhythmicity models in wheat (R2–R5) and artificial diet (R2–R4). Separate sheets provide the XSTREME results for each model.

**Table S11. Rhythmicity profiles of canonical insect circadian clock transcript homologs in *R. padi*.**

Excel file containing tables summarizing dryR and Nitecap rhythmicity results for *R. padi*transcript isoforms identified as homologs of canonical insect circadian clock genes. Results are shown for aphids feeding on wheat and artificial diet under light-dark (LD) and constant darkness (DD) conditions. The dryR classifications indicate inferred rhythmic models and associated parameters, and Nitecap results report transcript rhythmicity statistics and associated parameters for each condition.

## Notes

https://zenodo.org/records/23250260

### Competing Interest Statement

The authors have declared no competing interest.

