## Supplemental Note 1 for "Host Plant and Light Regime Shape Limited, Whole-Body Transcriptional Rhythmicity in the Bird Cherry-Oat Aphid (*Rhopalosiphum padi*)"

### Supplemental Note 1. Cross-method agreement in rhythmicity detection is limited and restricted to LD conditions

To assess the robustness of rhythmicity detection, dryR results were compared with those obtained using Cosinor and JTK_CYCLE, two commonly used algorithms for identifying rhythmicity in time-series data (Fig. S4; Table S5). Under LD conditions on wheat, Cosinor and JTK_CYCLE identified 3,350 and 1,053 transcripts, respectively, with significant 24 h rhythms (*τ* = 24 h, *p* < 0.05). Following FDR correction, 840 (Cosinor) and 265 (JTK_CYCLE) transcripts remained significantly rhythmic (*τ* = 24 h, *q* < 0.05). Under DD conditions on wheat, Cosinor and JTK_CYCLE detected 3,819 and 423 transcripts with significant 24 h rhythms (*τ* = 24 h, *p* < 0.05), respectively. However, after FDR correction, no transcripts remained rhythmic according to Cosinor, whereas 109 transcripts remained rhythmic using JTK_CYCLE (*τ* = 24 h, *q* < 0.05).

A similar pattern was observed for aphids on artificial diets, where rhythmic transcripts were detected primarily under LD conditions. With Cosinor and JTK_CYCLE, 2,496 and 453 transcripts were detected as having significant 24 h rhythmicity (*τ* = 24 h, *p* < 0.05), respectively. After FDR correction (*τ* = 24 h, *q* < 0.05), 372 and 77 transcripts remained rhythmic. In DD conditions on artificial diet, 1,877 (Cosinor) and 173 (JTK_CYCLE) transcripts exhibited nominally significant rhythms; however, no transcripts remained significant after FDR correction using either method (*τ* = 24 h, *q* < 0.05).

Overlap among all three methods was confined to LD conditions (Fig. S4A and C). Under LD on wheat, 138 rhythmic transcripts were jointly identified as rhythmic by all three approaches, corresponding to dryR models WhR2, WhR4, and WhR5. Under LD conditions on artificial diet, 50 transcripts were commonly detected as rhythmic by all three tools. In contrast, under DD conditions on both wheat and artificial diet, no transcripts were identified as rhythmic by all methods (Fig. S4B and D). However, in DD on wheat, 54 rhythmic transcripts were jointly identified in dryR (models WhR3, WhR4, WhR5) and JTK_CYCLE. Together, these comparisons reveal two trends. First, after FDR correction, all three methods show a general decline in detectable rhythmicity as cues are removed: light, host plant, or both. Second, at the nominal *p*-value threshold (*α* = 0.05), dryR and Cosinor detected more rhythmic transcripts in DD than LD on wheat, driven by the large WhR3 class; this exception was not reproduced by JTK_CYCLE and did not survive FDR correction, indicating these transcripts have a weaker or more noisy oscillatory structure (Fig. S4E). Additionally, cross-method agreement is strongest for LD-driven rhythms. In contrast, rhythms for diet-fed aphids and those reared under constant darkness are more sensitive to the choice of detection algorithm.
